# Collaborative Cross mice have diverse phenotypic responses to infection with Methicillin-resistant *Staphylococcus aureus* USA300

**DOI:** 10.1101/2023.07.12.548741

**Authors:** Aravindh Nagarajan, Kristin Scoggin, Jyotsana Gupta, Manuchehr Aminian, L. Garry Adams, Michael Kirby, David Threadgill, Helene Andrews-Polymenis

## Abstract

*Staphylococcus aureus* (*S. aureus*) is an opportunistic pathogen causing diseases ranging from mild skin infections to life threatening conditions, including endocarditis, pneumonia, and sepsis. To identify host genes modulating this host-pathogen interaction, we infected 25 Collaborative Cross (CC) mouse strains with methicillin-resistant *S. aureus* (MRSA) and monitored disease progression for seven days using a surgically implanted telemetry system. CC strains varied widely in their response to intravenous MRSA infection. We identified eight ‘susceptible’ CC strains with high bacterial load, tissue damage, and reduced survival. Among the surviving strains, six with minimal colonization were classified as ‘resistant’, while the remaining six tolerated higher organ colonization (‘tolerant’). The kidney was the most heavily colonized organ, but liver, spleen and lung colonization were better correlated with reduced survival. Resistant strains had higher pre-infection circulating neutrophils and lower post-infection tissue damage compared to susceptible and tolerant strains. We identified four CC strains with sexual dimorphism: all females survived the study period while all males met our euthanasia criteria earlier. In these CC strains, males had more baseline circulating monocytes and red blood cells. We identified several CC strains that may be useful as new models for endocarditis, myocarditis, pneumonia, and resistance to MRSA infection. Quantitative Trait Locus (QTL) analysis identified two significant loci, on Chromosomes 18 and 3, involved in early susceptibility and late survival after infection. We prioritized *Npc1* and *Ifi44l* genes as the strongest candidates influencing survival using variant analysis and mRNA expression data from kidneys within these intervals.

**Author Summary:** Methicillin-resistant *Staphylococcus aureus* is a human opportunistic pathogen that can cause life-threatening diseases. To study the influence of host genetics on the outcome of MRSA infection, we infected a collection of genetically diverse mice. We identified different phenotypes for survival, organ colonization, and tissue damage, and classified CC strains into MRSA susceptible, tolerant, and resistant categories. We identified several parameters that correlated with these phenotypes. Four CC strains exhibited strong sexual dimorphism in infection outcome: females lived longer, and males had higher baseline circulating monocytes and red blood cells. Several of the CC strains we characterize may represent better animal models for diseases caused by MRSA. QTL analysis identified two genes, *Npc1* and *Ifi44l*, as strong candidates for involvement in early susceptibility and late survival after MRSA infection. Our data suggests a strong involvement of host genetics in MRSA infection outcome.

## Introduction

*Staphylococcus aureus*, a gram-positive organism, is a commensal and opportunistic pathogen that can cause severe morbidity and mortality [1, 2]. This organism colonizes 20-30% of humans permanently and another 50-60% intermittently [3, 4]. *S. aureus* colonization is a risk factor for subsequent infections ranging from mild skin and soft tissue infections to serious invasive infections and death [5–7]. *S. aureus* is a multi-host pathogen that can colonize and cause disease in cows, chickens, pigs, sheep, and wild animals [8]. Methicillin-resistant *S. aureus* is resistant to β-lactam antibiotics [9], and was primarily associated with hospital-acquired infections until the 1980s [10, 11]. In the 1990s, MRSA spread rapidly in the community and is now considered a global threat [12]. MRSA causes more than 300,000 hospitalizations and 10,000 deaths in the US annually, at an estimated cost of 1.7 billion dollars (CDC, 2018).

USA300 is a highly successful community-acquired clone of MRSA. First reported in 1999, by 2011 USA300 became the primary cause of severe skin and soft-tissue infections, bacteremia, and community-onset pneumonia [13]. The increased disease burden caused by MRSA USA300 is attributed to the presence of multiple mobile genetic elements (MGEs) carrying virulence factors. In humans, USA300 infections are associated with increased mortality and increased incidence of severe sepsis than hospital-acquired MRSA infections [14, 15]. The differences in colonization rates, diversity in disease progression and mortality, exhibited by *S. aureus* suggests the involvement of host genetics.

Studies investigating the pathogenesis of *S. aureus* have relied heavily on a limited number of inbred mouse strains [16, 17]. *S. aureus* infections in mice induce a diverse spectrum of diseases similar to what is observed in humans [17]. Intravenous infection in mice with *S. aureus* is commonly used as a model for sepsis with this pathogen [17–20]. Following intravenous inoculation with up to 10^7^ organisms, *S. aureus* Newman (a methicillin-sensitive strain) eventually disseminates into various tissues, leading to the formation of abscesses in the vasculature, lung, heart, liver, spleen, and kidney [21]. It can take up to a month for these accumulating lesions to cause death in mice [21]. At higher doses (5×10^7^-5×10^8^ CFU), mice develop septic shock within 12-48 hours leading to death [22]. Infection with MRSA USA300 in mice follows a similar pattern of organ colonization [23].

Very little is known about the host genes and gender differences that influence *S. aureus* infections in mice. Targeted deletion of MyD88 (myeloid differentiation primary response protein, C57Bl6 x 129S1 intercross) and NOD2 (nucleotide-binding oligomerization domain-containing protein 2) render mice highly susceptible to *S. aureus* infection [24, 25]. Outcomes after infection with *S. aureus* differ depending on the strain of laboratory mice used. In a bacteremia model, C57BL/6J mice are the most resistant in controlling bacterial growth and survive, while A/J, DBA/2, and BALB/c mice are very susceptible [26]. This information has been used to map a limited number of loci involved in this differential outcome. Using C57BL/6J and A/J Chromosomal substitution strains (CSS), six genes on three different chromosomes (Chr 11 – *Dusp3*, *Psme3*; Chr 8 – *Crif1, Cd97*; Chr 18 – *Tnfaip8, Seh1l*) have been implicated to susceptibility after infection with *S. aureus* (sanger 476) strain [27–29]. To our knowledge, no study has looked at the host factors involved in USA300 pathogenesis, a hypervirulent isolate compared to other strains [30–32]. Furthermore, no screen for MRSA infection disease outcomes has been performed across a broad range of host genetics.

Collaborative Cross mice have been successfully used to create disease models and identify susceptibility genes for various infections [33]. The CC is a large panel of recombinant inbred mouse strains created by interbreeding five classical inbred strains (A/J, C57BL/6J, 129S1/SvlmJ, NOD/ShiLtJ, NZO/HlLtJ) and three wild-derived inbred strains (CAST/EiJ, PWK/PhJ, and WSB/EiJ) using a funnel-breeding scheme [34, 35]. CC strains have more than 30 million Single Nucleotide Polymorphisms (SNPs), and their genomes are fully sequenced, making them an ideal population for QTL mapping [36, 37]. The CC has been heavily used to identify susceptibility to viral pathogens, including Ebola virus [38], West Nile virus [39], Influenza A virus [40], Coronaviruses [41, 42], Theiler’s murine encephalomyelitis virus [43], Cytomegalovirus [44] and Rift Valley fever virus [45]. The CC has also been used to study infection with bacterial pathogens, including *Mycobacterium tuberculosis* [46, 47], *Klebsiella pneumonia* [48], *Pseudomonas aeruginosa* [49], *Borrelia recurrentis* [50], *Salmonella* Typhimurium [51, 52] and *Salmonella* Typhi [53].

We were interested in how host genetics influences disease outcome after MRSA bacteremia. Therefore, we screened 25 CC strains for disease outcome phenotypes after intravenous infection with MRSA USA300. We chose a 7-day screening period and very sensitive telemetry monitoring of MRSA infected animals in an attempt to capture and identify range of disease phenotypes from highly susceptible to resistant. Using this model system, we identified a wide range of disease outcomes, potential new MRSA disease models, and genomic regions linked to survival using QTL mapping. In order to prioritize the genes within these intervals we used variant analysis and mRNA expression data from kidneys.

## Results

### CC strains have diverse disease outcomes after MRSA infection

We screened twenty-five Collaborative Cross strains (three males and three females per strain) for their phenotypes after intravenous infection with MRSA USA300 (Table S1). Because blood borne infections can become rapidly fatal, we used a 7-day screening period to allow us to delineate sensitive and resistant CC lines. Animals were monitored for clinical symptoms using a continuously monitored telemetry system and a manual health scoring method up to one-week post-infection to identify those susceptible to MRSA infection. At the time of necropsy spleen, liver, heart, lung, and kidney were collected to enumerate the bacterial load and tissue damage (Table S2).

C57BL/6J mice, known to survive MRSA infection, served as controls [26]. All C57BL/6J mice survived until the end of the experiment (day 7) and were thus classified as a surviving strain (Fig 1A). CC strains varied widely in their response to MRSA sepsis (Fig 1 and Fig S1). Median survival ranged from 2.5 to 7 days (Fig 1A). Median weight loss ranged from losing 20% of their body weight (CC002, CC037) to losing less than 1% (CC003, CC017, CC024) (Fig 1B). Out of the 156 infected mice, 10 mice belonging to 6 different strains appeared to clear MRSA from all organs we collected (Fig 1D). We included these mice in our analysis based on disturbances in circadian pattern following infection and histology scores, suggesting that they did become infected (Fig S5 and S6).

**Fig 1:**
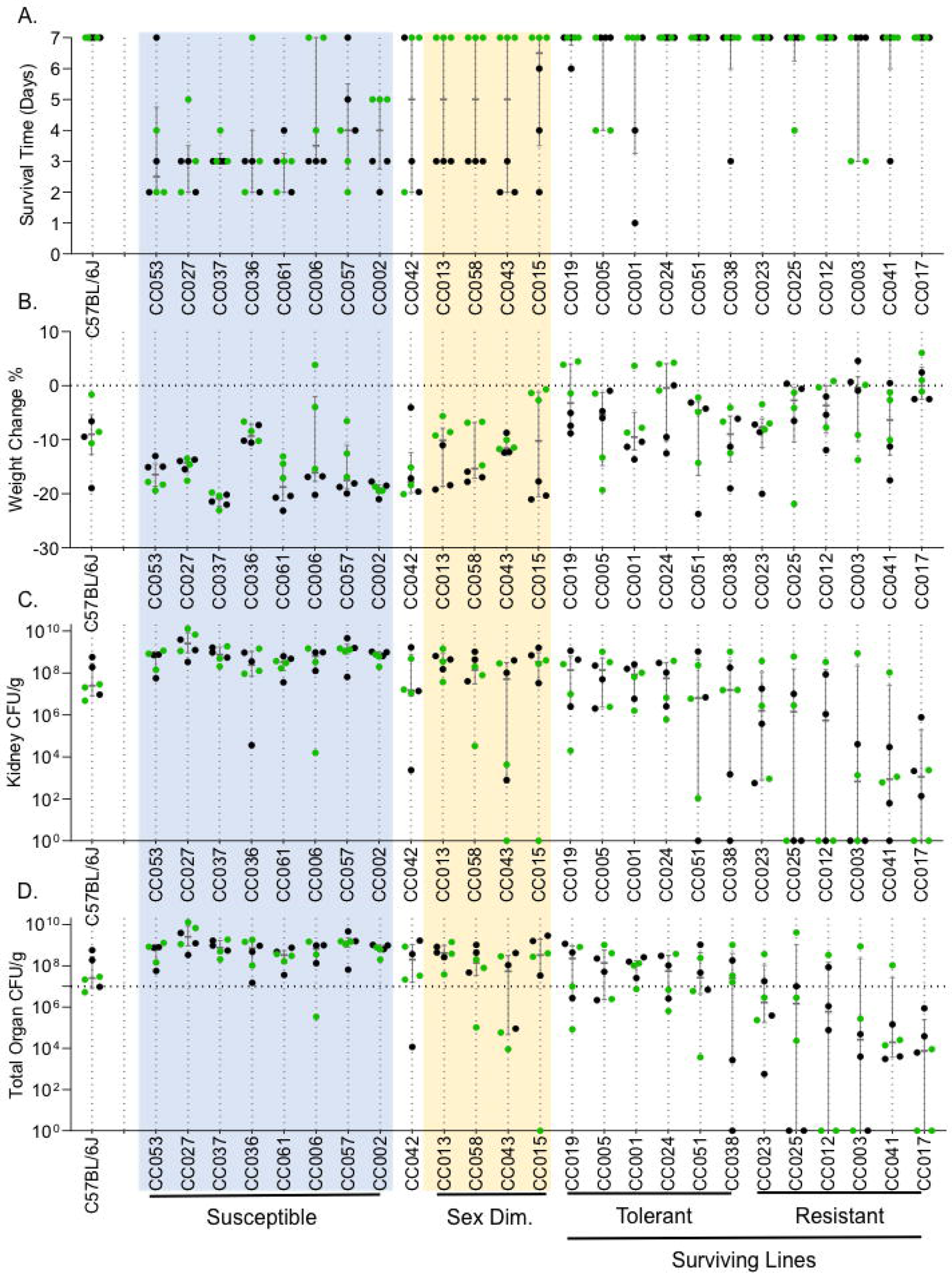
CC strains exhibit a diverse response to MRSA infection. After intravenous infection with MRSA USA300 as described in the Materials and Methods section, we show: A. survival time, B. percent weight change after infection, C. kidney colonization, and D. Total organ colonization; horizontal dotted line represents inoculum dose. CC strains are shown in the ascending order of survival. If survival was equal between two or more strains, strains are arranged in descending order of total organ colonization (the combined colonization of all organs collected). Dots represent individual mice; black dots represent males; green dots represent females. The median and interquartile range are shown for each strain.

CC strains that had four or more mice of the six infected meeting our euthanasia criteria before day seven were classified as ‘susceptible’. Eight of the 25 CC strains we infected were classified in this category (CC002, CC006, CC027, CC037, CC036, CC053, CC057, CC061) (Fig 1A). Susceptible strains, on average, lost 16% of their body weight (Fig 1B). Two susceptible strains, CC027 and CC057, had median kidney colonization on the order of 10^9^, ∼100-fold higher than the inoculation dose (Fig 1C).

Five CC strains had three mice that met our euthanasia criteria before day 7. Of these five strains, the survival of four of these strains was uniformly sexually dimorphic during our experimental time period: all the males met our euthanasia criteria early in the experimental period, while all females of these strains survived the full 7-day infection (CC013, CC015, CC043, CC058) (Fig 1). These four CC strains were classified as sexually dimorphic in their survival after MRSA infection. CC001, classified as ‘surviving’, may also display sexual dimorphism; 2/3 males met our euthanasia criteria prior to day 7, and the remaining male never recovered to a pre-infection circadian pattern of body temperature or activity. Infection experiments of longer duration will be required to determine if the sensitivity of males in this strain is uniform. Finally, although CC042 is highly susceptible to other bacterial pathogens [46, 51, 54], couldn’t easily be classified with respect to MRSA infection outcome as two females and one male survived infection.

We classified the remaining 12 surviving CC strains into tolerant and resistant phenotypes. We defined tolerance as surviving the infection with a high bacterial burden while limiting the health impact caused by the pathogen [55–57]. Six strains with higher total organ colonization compared to the inoculum dose were classified as tolerant (CC001, CC005, CC019, CC024, CC038, CC051) (Fig 1D). These strains limited the impact to their health (based on circadian pattern and health scores) despite high colonization in several organs (Fig S5 and Table S2). Although CC051 mice infected with MRSA survived and were classified here as tolerant, they exhibited some weight loss and a persistent disruption of their circadian pattern of body temperature and activity. Infections of longer duration will be needed to determine whether CC051 mice are truly tolerant or display a delayed susceptibility phenotype [52].

Surviving the infection by preventing colonization or clearing the pathogen was defined as resistance [55–57]. The remaining six strains, with lower (compared to the inoculum dose) or no colonization, were classified as resistant (CC003, CC012, CC017, CC023, CC025, CC041) (Fig 1D). MRSA infection had minimal impact on the health of these CC strains (Fig S5 and Table S2). The wide range of disease severity, survival, weight loss, and organ colonization we observed during blood borne MRSA infection suggest diverse host responses across genetically different CC strains.

### Correlation of colonization, tissue damage and immune parameters with survival

Pre-infection weight did not influence survival (R = −0.05), while weight loss after infection correlated with poor survival (R = −0.65) (Fig 2A). Colonization in all five organs was highly correlated with poor survival (Fig 2A). The inability to limit bacterial growth in the kidney has previously been associated with susceptibility to intravenous *S. aureus* infections in mice [26]. In our experiments, the kidney was the most highly colonized organ across all CC strains and high kidney colonization correlated with poor survival (R = −0.58) (Fig 1C, 2A). Unexpectedly however, colonization in the liver (R = −0.78), spleen (R = −0.73), and lung (R = −0.67) was more highly correlated with poor survival than colonization of the kidneys (Fig 2A), despite the fact that these organs had lower bacterial load than the kidneys (Fig 1C, 1D) and less tissue damage (Fig 2B). These data suggest that the spleen, liver and lung are more intolerant to colonization by MRSA and acute damage than the kidney during MRSA infection.

**Fig 2:**
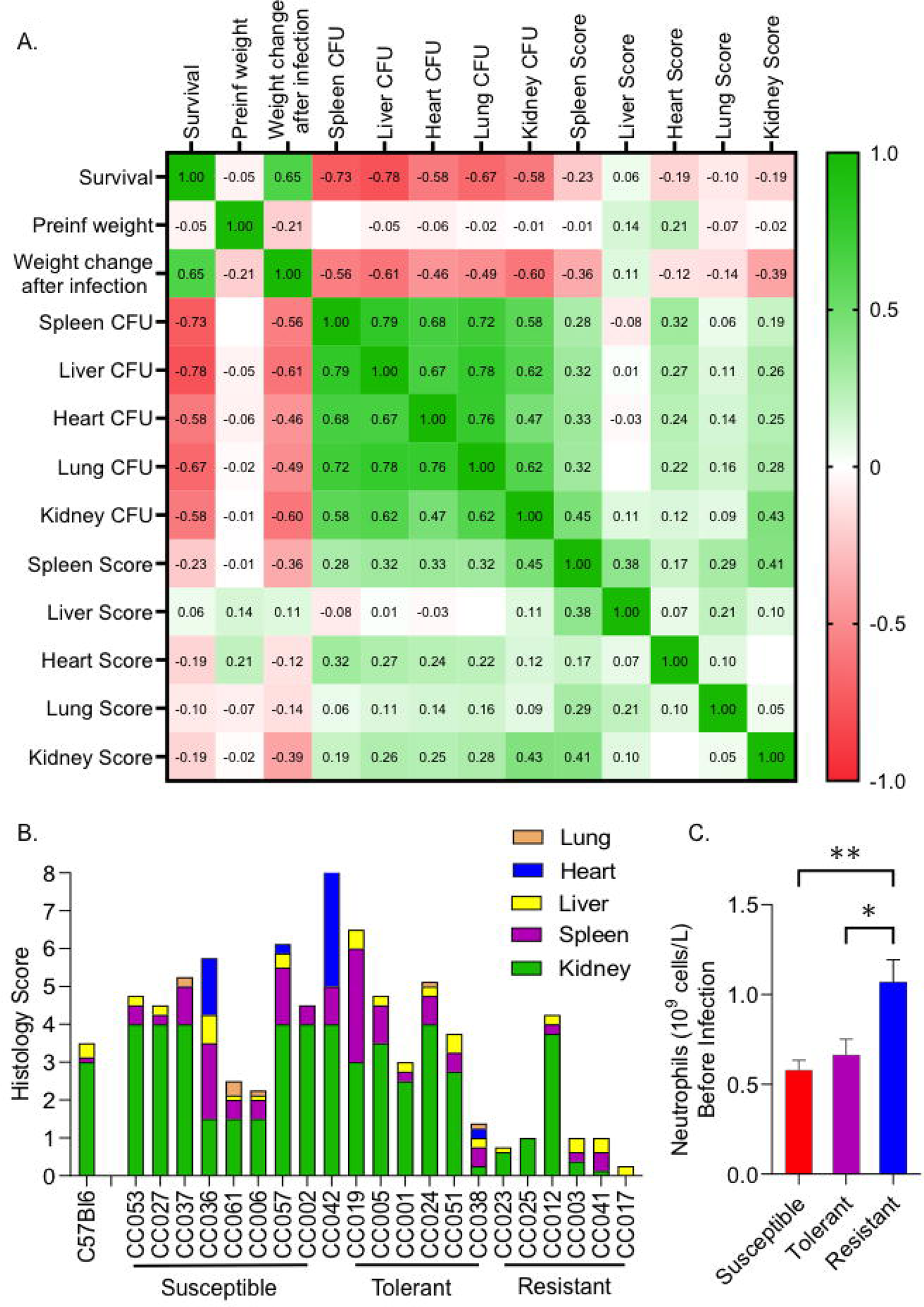
Correlation of tissue colonization, histology scores and blood parameters with survival. A. Heat map showing Spearman correlation ‘R’ values between survival, weight change, organ colonization, and tissue damage scores. B. A stacked bar plot showing the median tissue damage score for five organs (Kidney = green, spleen = purple, liver = yellow, heart = blue, lung = orange). Pathology in each tissue was blindly scored by a pathologist on a scale of 0 = no damage, to 4= severe damage. C. Mean neutrophil levels in uninfected mice. Mann-Whitney tests were performed to determine statistical significance (* = P < 0.05, ** = P < 0.01).

*S. aureus* causes damage to host tissue in mice as early as 48 hours after intravenous challenge [21]. To assess tissue damage in the CC mouse population, a board-certified pathologist scored each tissue blindly for damage (0 = normal to 4 = severe damage) (Table S3 and Fig S2). Across all organs, resistant strains had significantly less tissue damage than tolerant (P < 0. 001) and susceptible strains (P < 0.010) (Fig 2B). Interestingly, there was no significant difference in tissue damage between tolerant and susceptible strains (P = 0.57) (Fig 2B). This finding suggests that tolerant strains may survive MRSA infection in the face of both high colonization and severe tissue damage.

High bacterial burden in kidneys translated to more significant tissue damage. Of the 25 CC strains we studied, 16 had a median pathology score of 2.5 or above for kidneys, indicating moderate to severe tissue damage (Fig 2B, Scale of 0-4, 0= no damage, 4= severe damage). This damage did not translate to susceptibility however, as histology scores across all five tissues that we sampled had only a weak correlation with poor survival (Fig 2A). Spleen damage had the highest correlation R = −0.23 with poor survival, followed by kidney (R = −0.18) and heart (R = −0.18) damage. This weak correlation may be due to the diverse phenotypic responses exhibited by the CC strains to MRSA infection. Some infected CC strains met our euthanasia criteria very early in infection, perhaps before extensive tissue damage had time to occur. Other CC strains lived to the end of the experiment, allowing time for severe tissue damage to occur and become cumulative.

Pre-infection inflammation and immune parameters play a role in disease prediction and progression [42, 58]. We collected blood from mice five weeks before infection to perform Complete Blood Counts (CBC). Interestingly, CC strains resistant to MRSA infection had a higher number of pre-infection circulating neutrophils than susceptible and tolerant strains (Fig 2C). There were no significant differences in other pre-infection CBC parameters (Table S2).

### Survival is sex-dependent in four CC strains

In the four sexually dimorphic strains, females lived until the end of the experiment (day 7), but all the males met our euthanasia criteria much earlier (Fig 3A). Males lost more weight than females (Fig 3B) and had higher liver, spleen, and lung colonization (Fig 3C). Colonization did not differ significantly between males and females in the kidneys, the highest colonized organ in these strains (Fig 3C). Females of these CC strains appear to limit MRSA replication in some organs or spread from the kidneys by unknown mechanisms.

**Fig 3:**
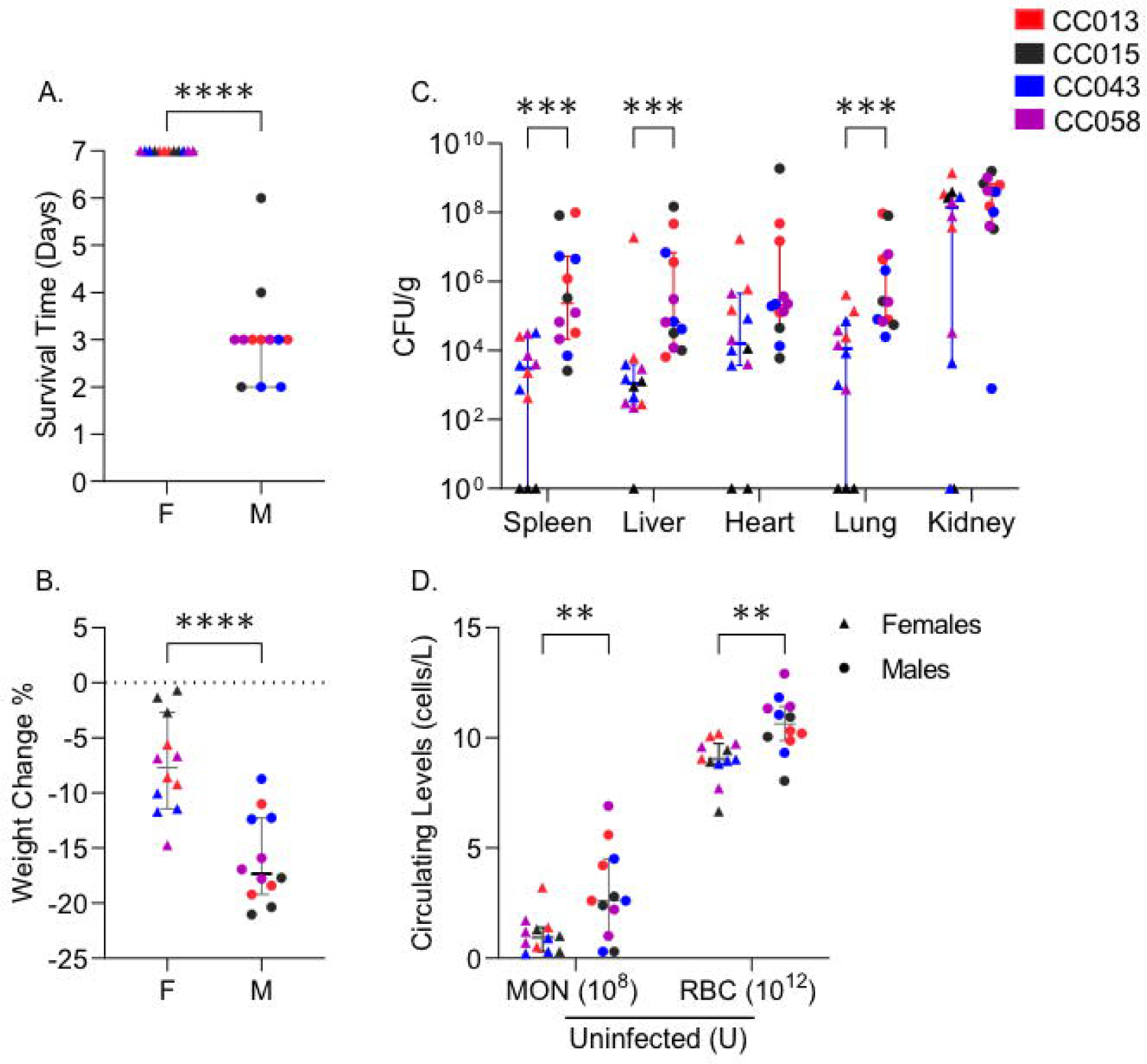
Sexual dimorphism in survival after MRSA infection. We present pooled data for the four strains that exhibited sexual dimorphism in survival: A. survival time, B. percent weight change after infection, C. colonization across organs, and D. pre-infection monocyte and red blood cell counts (U – Uninfected). Each point represents an individual mouse; filled circles represent males; triangles represent females. Each color represents a CC strain as noted in the legend. Medians with 95% confidence interval are shown for each strain. A, B, and D – Student’s T-test and C

To assess baseline differences between males and females of the strains that were sexually dimorphic with respect to MRSA infection, we evaluated pre-infection complete blood counts (CBC). Males had higher circulating monocytes and red blood cells before infection than females of these strains (Fig 3D). The number of circulating monocytes (P = 0.83) and red blood cells (P = 0.10) were not significantly different between males and females in the rest of the CC population (21 strains) (Table S2). Thus, this difference in the starting blood parameters is unique to CC strains that displayed sexual dimorphism in susceptibility to MRSA infection.

### Baseline body temperature and baseline activity did not correlate with survival

To track the disease progression in infected mice, we used a combination of automated telemetry and manual health scoring (Table S2). A surgically implanted telemetry device tracked body temperature and activity levels every minute before and throughout the infection (Fig S5 and Fig S6). Data from five days before infection was used to create a baseline circadian pattern for each mouse. A machine-learning algorithm was also used to predict when infected mice deviated from their normal circadian pattern after infection. No significant differences were identified in the elapsed time between infection and deviation from the normal circadian pattern between surviving and susceptible strains (Table S2). 90% of the infected mice deviated from their normal circadian pattern of body temperature and activity within 24 hours of infection (Fig S3). The first health check performed by laboratory staff, 24 hours after infection, could only detect 56% of these mice. In the next two days, twice daily health checks detected all the remaining mice as ill (Fig S3). These data suggest a significant lag time between circadian pattern disruption detected by telemetry and signs of illness detected by laboratory staff and the effectiveness of the telemetry system in accurately predicting survival after infection.

### Two independent genomic regions influence survival

We used RQTL2 software with CC genotypes imputed from qtl2qpi to identify genomic regions associated with survival after MRSA infection [59, 60]. Within the CC strains we infected, survival was a highly heritable trait with a broad sense heritability score of 0.38 (Table S4). While using only 25 CC strains reduces the power of QTL mapping [61], using 6 animals per strain and continuous monitoring using a telemetry device added more power to our study and allowed us to sensitively define the phenotypes we observed.

For early susceptibility on day 2, there was a highly significant peak on chromosome 18 (Fig 4A). We named this QTL peak ESMI (Early Susceptibility in MRSA Infection). This peak disappeared after day 3 post-infection. For survival on day 7 (late survival), there was a highly significant peak on Chromosome 3 (Fig 4B). We named this peak LSMI (Late Survival in MRSA Infection). To understand the impact of sex on survival, we performed the same analysis after removing the four strains that exhibited sexual dimorphism in survival. The early susceptibility peak on Chr 18 and late survival peak on Chr 3 were retained (Fig S4), suggesting that sex did not influence our QTL peaks. We compared the phenotype at the top SNPs in both peaks for potential interaction (Table S5). No significant interactions were identified, suggesting that our QTL regions act independently of each other.

**Fig 4.**
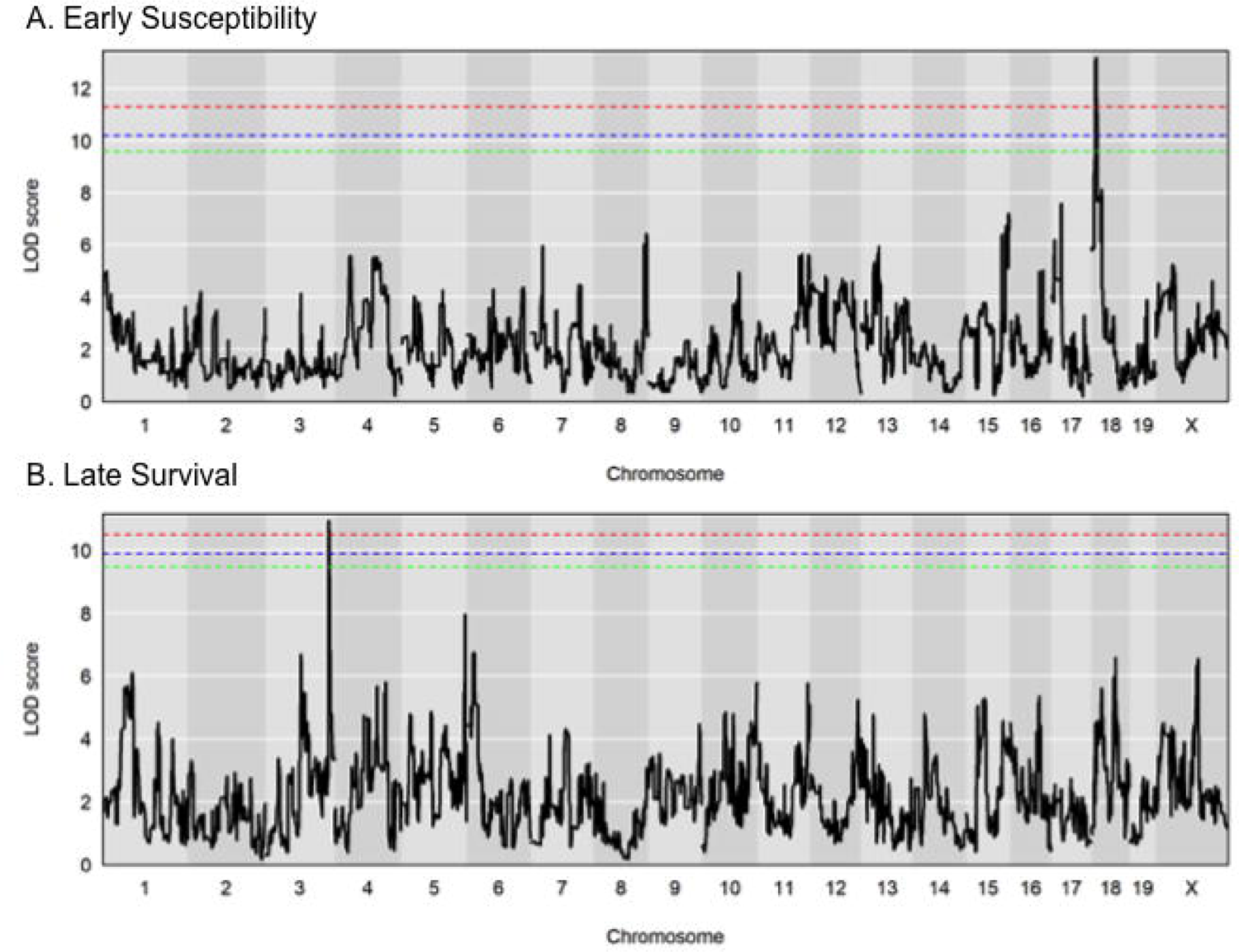
Genomic regions involved in survival after MRSA infection. LOD plots for square root transformed survival phenotype (percentage of animals that survived at the end of each day). A. Early susceptibility in MRSA infection (ESMI, day 2 post-infection) peak on chromosome 18. B. Late survival in MRSA infection (LSMI, day 7 post-infection) peak on chromosome 3. The dotted (Red – 95%, Blue – 90%, Green – 85%) lines represent the significant LOD scores for 999 permutations.

### *Npc1* is a strong candidate for influencing early susceptibility during MRSA infection

The advantage of the CC panel over other outbred mouse panels is the high number of SNPs and structural variants present across the CC genomes [36, 37]. We used this feature to shortlist the genes within the associated genomic regions to identify those of most interest. The early susceptibility QTL peak, ESMI, on chromosome 18 is a 5 Mb region between 12-17 Mb with a peak around 14.67 Mb. This region contains 21 protein-coding genes and 49 non-coding RNA genes. Because the CC strains are a mosaic of eight inbred founder strains, we looked at the founder effect plot in this region to attempt to identify each founder’s contribution to survival (Table S5). We found that the WSB allele in this region contributes to reduced survival, while A/J and NZO alleles contribute to prolonged survival (Fig 5A). Using all the SNPs, insertions and deletions that matched this allele effect pattern in the Mouse Phenome Database (MPD), we identified 20,582 variants across all 21 genes. We used Variant Effect Predictor (VEP) to further prioritize these variants based on their predicted effect on protein structure and function, identifying 52 high and moderate-impact variants in 7 genes (Table S6). The highest impact variant for each gene is listed (Fig 5B). Four of the seven genes we shortlisted have previously been associated with immune function in the Mouse Genome Informatics Portal (highlighted in red, Fig 5B) [62]. We sequenced mRNA from kidneys collected from CC strains at necropsy (at least one male and one female per strain) to further prioritize genes. Then, we looked for differentially expressed genes that matched the founder effect pattern within this region. For the early susceptibility QTL peak, two genes, Cadherin2 (*Cdh2*) and NPC intracellular cholesterol transporter 1 (*Npc1*), matched the founder effect pattern (Fig 5C). *Npc1* is our top candidate in this region due to its previous association with immune functions [63, 64].

**Fig 5.**
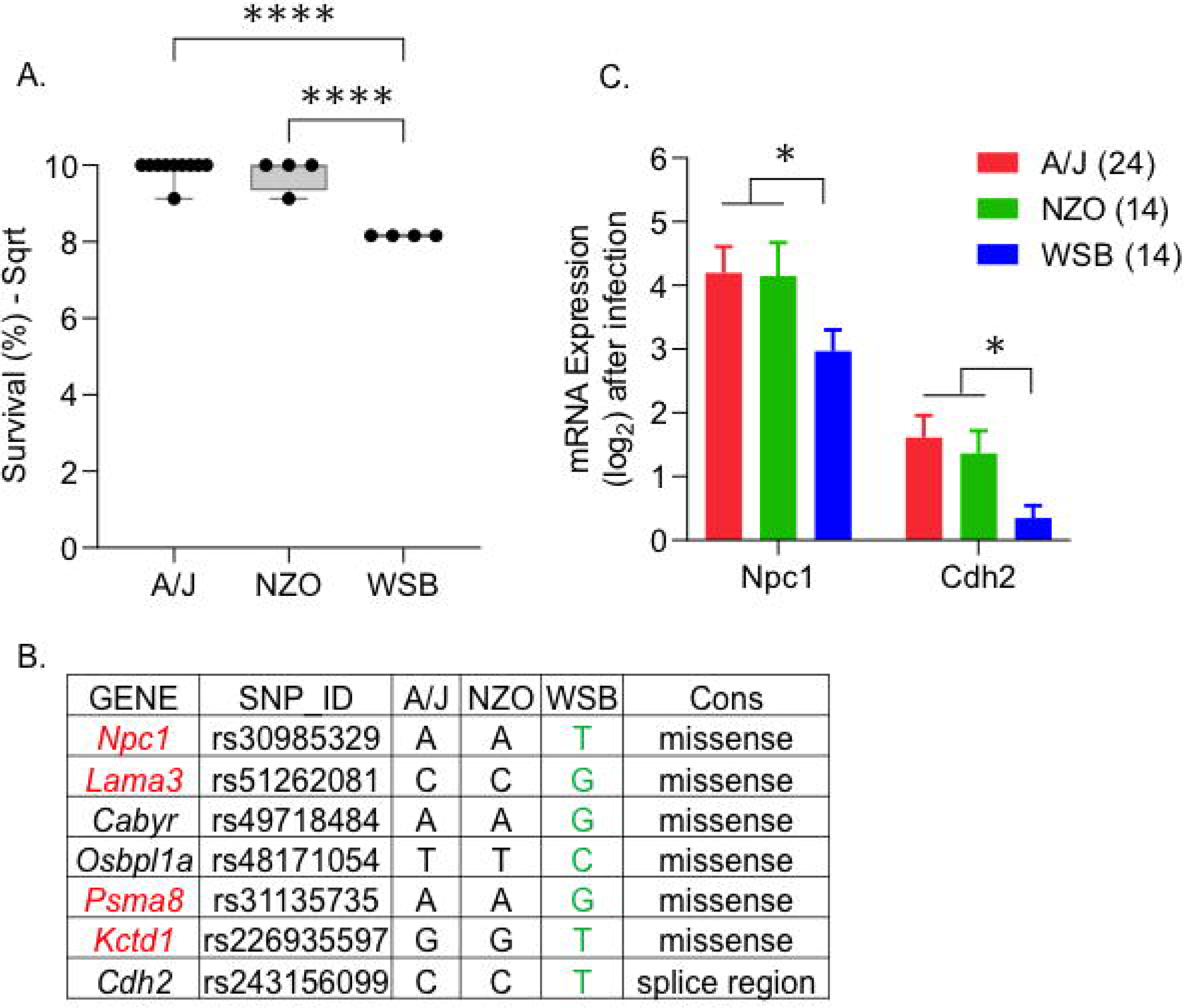
Strongest candidate genes on chromosome 18 for early susceptibility. A. Box plot showing the founder allele contribution at the highest marker on the peak. Each dot represents a strain, lines represent minimum to maximum. B. Genes with high-impact variants shortlisted using the founder effect pattern. C. Mean kidney mRNA expression values after infection (log-transformed). The number of animals is given within brackets. Lines represent standard error of the mean values. ANOVA with multiple comparison correction using Tukey test was performed (**** = P < 0.0001).

### *Ifi44l* is a strong candidate for involvement in late survival after MRSA infection

The late survival QTL peak, LSMI, on chromosome 3 is a 4 Mb region between 146-150 Mb with a peak near 146.66 Mb. This region contains 13 protein-coding genes and 39 non-coding RNA genes. The A/J allele in this region reduced survival significantly, while CAST, NOD, NZO, and WSB alleles promoted longer survival (Fig 6A). Using this founder effect pattern, we shortlisted five genes with variants predicted to have a high impact on the function of the encoded proteins (Table S6). The top variant for each gene is listed (Fig 6B). Two genes in the Interferon Stimulated Gene family (ISG), Interferon induced proteins 44 and 44l genes (*Ifi44* and *Ifi44l),* are our primary candidates at this locus. *Ifi44* has a missense mutation, and *Ifi44l* has a high-impact mutation in the gene’s promoter region. In mice and humans, *Ifi44l* is a paralog of *Ifi44, but it remains* unclear if *Ifi44* and *Ifi44l* have redundant or distinct functions [65]. Using gene expression data, we determined that *Ifi44l*, is highly expressed in strains with the A/J allele but not with the alleles from the other founders (Fig 6C). Thus, *Ifi44l* became our top candidate gene influencing late survival during MRSA infection.

**Fig 6.**
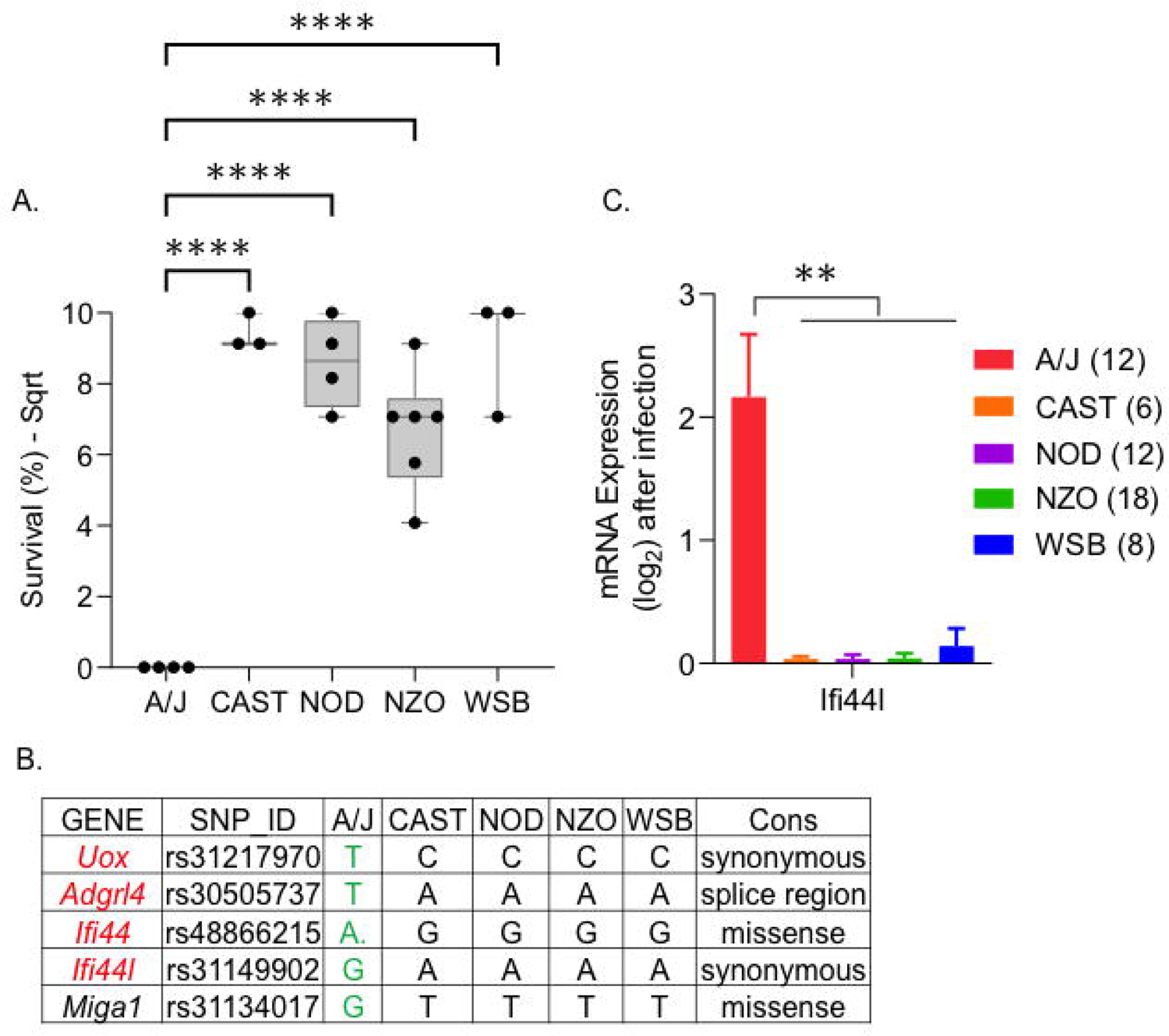
Strongest candidate genes on chromosome 3 for late survival. A. Box plot showing the founder allele contribution at the highest marker on the peak. Each dot represents a strain, lines represent minimum to maximum. B. Genes with high-impact variants shortlisted using the founder effect pattern. C. Mean kidney mRNA expression values after infection (log-transformed). The number of animals sequenced for each founder allele is within the brackets. Lines represent standard error of the mean values. ANOVA with multiple comparison correction Tukey test was performed (**** = P <0.0001).

## Discussion

We used a genetic screen of CC mouse strains to determine whether mice of different genetic backgrounds have diverse disease outcomes after intravenous infection with MRSA USA300, and ultimately to identify regions of the genome linked to survival. We identified 8 CC strains that were highly susceptible to fatal infection, 4 strains with strong sexually dimorphic survival, 6 strains that appeared tolerant, and 6 strains that were resistant to MRSA infection (even totally clearing the pathogen in several cases). Survival, bacterial colonization, and weight loss differed even within a given phenotypic category. Across 8 CC strains that we determined to be susceptible to MRSA sepsis based on reduced survival after infection, survival time ranged from 2.5 to 4 days, and kidney colonization varied as much as 10-fold. Two susceptible strains, CC061 and CC036 had very similar kidney colonization, yet CC061 appeared to have a more severe disease and lost twice as much weight as CC036 (19% vs. 9%). We also observed similar diversity in weight loss and colonization within tolerant and resistant strains. Our data support both the importance of host genetics in disease outcome after MRSA infection, but also the hypothesis that there may be multiple mechanisms of susceptibility, tolerance and resistance to MRSA infection across a broad set of host genetics.

In both animals and humans, males are more susceptible than females to bacterial infections [66]. Some of these differences can be attributed to the differences in X-chromosome number and inactivation process between males and females. The X-chromosome also encodes several critical immune genes that may play important roles in sexually dimorphic infection outcomes [67]. Males have a higher overall risk for MRSA carriage [68] and bloodstream infections [69, 70]. However, there is no consensus on the role of gender on mortality [70–74]. Across all CC strains in our study females survived longer than males. We determined that this difference was driven by 4 CC strains that exhibited strong sexual dimorphism in survival, despite similar levels of kidney colonization in both sexes.

Two blood parameters, pre-infection baseline numbers of circulating monocytes and red blood cells, were higher only in males of the CC strains that exhibited sexually dimorphic survival phenotypes in response to MRSA infection. This finding suggests that increased numbers of circulating monocytes and red blood cells may be linked to susceptibility of males of these CC strains to severe MRSA infection. *S. aureus* persists within monocyte-derived macrophages and can use these cells as a vehicle for dissemination [75, 76]. Furthermore, heme iron is the preferred source of iron for *S. aureus* during infection [77], and this organism can bind and lyse RBCs [78–80] and directly bind hemoglobin [81] [82]. Thus, in males of these CC strains, increased pre-infection circulating monocytes may facilitate more efficient MRSA dissemination while increased heme availability may provide an additional source of iron for MRSA during infection.

Neutrophils are the primary defense against *S. aureus* infections [7, 83], and patients with impaired neutrophil numbers and/or function are predisposed to *S. aureus* infections [84–86]. We noticed that pre-infection circulating neutrophil numbers were significantly higher in our MRSA-resistant CC strains than in the tolerant and susceptible strains. Our data is consistent with observations in humans, supporting a critical role for neutrophils in resistance to MRSA infection in several CC strains.

Most of the CC strains we used in this study were also used to define disease outcome after oral infection with non-typhoidal *Salmonella* [51, 52]. Six of the eight CC strains that were susceptible to MRSA infection were also susceptible or delayed susceptible to *Salmonella* infection (CC006, CC027, CC036, CC037, CC053, CC061). The remaining two MRSA susceptible CC strains, CC002 and CC057, were resistant and tolerant to *Salmonella* infection respectively [52]. One CC strain, CC038, was tolerant to both MRSA and *Salmonella* infection. However, all of the remaining CC strains we tested had different disease outcomes in MRSA infection than they did after *Salmonella* infection. The phenotypes we discovered suggest that some of the mechanisms of susceptibility to severe MRSA and *Salmonella* infection may be shared, while the mechanisms of tolerance and resistance to these two pathogens are largely distinct.

We compared our MRSA infection outcome data to a similar study of *Mycobacterium tuberculosis* (Mtb) infection in CC strains [47]. While this study did not classify survival phenotypes, we used the median lung colonization values as a rough proxy for disease severity. For the 52 CC strains infected with Mtb, log lung CFU ranged from 5.1 to 8.5, with a median of 6.7 CFU [47]. Among the six MRSA-susceptible CC strains that were also infected in *Mtb* studies, two strains (CC027, CC037) were sensitive to both *Salmonella* and MRSA infection and were also heavily colonized by *Mtb* in the lung [47, 51, 52]. A common mechanism may underlie the susceptibility of these CC strains to three very different bacterial pathogens, administered by different routes.

We also identified CC strains that were not uniformly susceptible to all three pathogens. Two CC strains susceptible to both MRSA and *Salmonella* (CC006, CC061) were poorly colonized in the lung with *Mtb* (5.6 and 6.5 log CFU) [47]. Finally, all five MRSA-resistant CC strains were highly colonized in the lung with *Mtb* (average – 8.4 log CFU), and thus appear to be potentially highly susceptible to *Mtb* infection. The MRSA-tolerant strains had variable *Mtb* colonization in the lung. This differential response to different bacterial pathogens suggests that most CC strains that we tested do not have a significant general dysfunction of immunity, where we would expect broad susceptibility to multiple pathogens. Future work will tease out specific mechanisms of susceptibility and resistance to different pathogens.

Using the survival data that we generated in this study, we identified QTL linked to both early susceptibility and late survival after intravenous MRSA infection. The primary candidate gene in the early susceptibility QTL, *Npc1*, is in endosomal and lysosomal membranes and mediates intracellular cholesterol trafficking [87]. In humans, mutations in *NPC1* cause Niemann–Pick Type C disease, leading to massive accumulation of cholesterol in lysosomes in all tissues and premature death [88]. Defects in *NPC1* can lead to increased lipid storage in macrophages and a chronic inflammatory condition [89, 90]. *NPC1* is also the primary host factor responsible for Ebola, and other filovirus, entry into host cells [91]. These viruses use *NPC1* on endosomes and lysosomes to trigger a membrane fusion process that allows the expulsion of the viral genome into the cytoplasm and subsequent viral replication [92].

Little is known about the role of *NPC1* in bacterial infections. In a hematopoietic mouse model, *Npc1* mutant mice have a significant increase in the relative abundance of *Staphylococcus* ssp. in their intestine [93], suggesting a role for *Npc1* in keeping *Staphylococcus* ssp. at a low level in this niche. Mutations in *Npc1* impair steady-state autophagosome maturation and interfere with the autophagic trafficking of bacteria to lysosomes in macrophages [94]. *S. aureus* manipulates autophagy in both neutrophils and macrophages to promote its survival and escape [95–97]. CC strains carrying the WSB allele in this region have a missense mutation in *Npc1*. This altered form of *Npc1* may interfere with autophagy in macrophages and neutrophils, such that *S. aureus* can survive initial phagocytosis leading to the early susceptibility phenotype.

The A/J allele on Chromosome 18 contributed to poor survival of MRSA infected mice on day 7. A/J mice are also highly susceptible to SH1000, a different strain of *S. aureus* [26]. *Ifi44l* promotes macrophage differentiation during bacterial infection and facilitates inflammatory cytokine secretion [98]. Overexpression of *Ifi44* or *Ifi44l* is sufficient to control respiratory syncytial virus (RSV) infection [65]. *Ifi44l* represses the interferon response, protecting the host from the negative effects of the innate immune response during viral infection [99]. The A/J *Ifi44l* allele may be less effective in controlling the interferon response, reducing the survival of *S. aureus* infected mice in the later stages of infection.

A series of previous studies examined the genetic differences that influence *S. aureus* Sanger-476 infection in A/J and C57BL/6J mice using a chromosome substitution strain (CSS) model [27–29]. Although C57BL/6 and A/J mice are among the CC founders, we did not identify any genes in common with this previous work. Like USA300, Sanger-476 is a community-acquired strain but is sensitive to methicillin [100]. USA300 also has additional pathogenicity islands that are absent in the Sanger-476 strain [101] and also produces higher amounts of virulence factors compared to other MRSA strains [30, 102, 103]. These genetic differences make USA300 distinct from, and potentially more virulent than, other community-acquired clones of MRSA. On the host side, the CC founders include six additional inbred strains, making the CC strain collection considerably more genetically diverse than the CSS panel. The fact that the genetic regions we identify as linked to survival in this study did not overlap with those of the previous study is thus likely due to use of different *S. aureus* strains, more diverse mouse strains and different mapping power [27]. However, given that earlier work using the CSS panel identified increased inflammation and defective immune responses as critical to MRSA resistance, we were encouraged to identify *Npc1* and *Ifi44l* as potential influencers of survival, as these genes are also involved in modulating inflammation and influencing immune responses.

The CC panel has previously been used to identify mouse strains for new model generation [33]. Although it was not a goal of this study to identify new mouse models for different MRSA caused syndromes, our analysis of pathology across different CC strains suggested that several strains may be make suitable models to fill gaps in our knowledge of cardiac disease, pneumonia, and recovery from *S. aureus* systemic infections. *S. aureus* is the leading cause of infective endocarditis (IE), which is associated with high mortality (40-50%) [104, 105]. Current models of IE require anesthetizing C57BL6/J mice and physically damaging their heart valve using a catheter [106, 107]. *S. aureus* is also the most common cause of bacterial myocarditis, with fatal outcomes [108–110]. Currently, there are no murine models to study MRSA induced myocarditis. In our study, three CC strains (CC013, CC036, C0042) had over 200-fold greater colonization in the heart than C57BL/6J mice and had systemic embolic bacterial purulent myocarditis. Similarly, pneumonia caused by *S. aureus* has a high mortality rate [111]. In our study, CC013, CC058, and CC061, had significant lung damage. Finally, we identified six MRSA-resistant CC strains (CC023, CC025, CC012, CC003, CC041, and CC017) that robustly survive systemic infection, have lower bacterial colonization, and in most cases less tissue damage than the traditional resistant mouse model for MRSA: C57BL/6J mice [26, 29]. We hope that the lines we list above will be studied in more detail and may provide a useful resource for various *S. aureus* based syndromes.

In summary, using colonization, telemetry, survival, and histology data, we have identified diverse disease outcomes after MRSA USA300 infection in and described two novel genomic regions that influence survival at different stages after systemic infection. In addition, our work suggests several interesting pre-infection correlates that appear to influence survival after MRSA infection, and a connection between sex and susceptibility to MRSA infection in certain genetic backgrounds. With several *S. aureus* vaccines failing at human clinical trials [112, 113], our data support strong consideration of host genetics as an important factor while designing therapeutics and vaccines against pathogens. Finally, as the CC strains have now been infected with multiple bacterial pathogens, future will work will use the phenotypic diversity of host responses to infection of this mouse collection to define different mechanisms of susceptibility, tolerance and resistance.

## Acknowledgements

We thank Dr. Magnus Hook for providing the bacterial strain used in this study. We also thank Dr. Karl Broman (University of Wisconsin) for helpful discussions and troubleshooting problems in the rqtl2 google forum. We also would like to thank Dr. Callie Kobayashi, Dr. Joana Rocha, Ms. Marissa Talamantes, Mr. Chris Bowden, Mr. Connor Mathis, and Ms. Kaya Mariello for assistance with experiments. This work was funded by the Defense Advanced Research Project Agency (DARPA), project DARPA D17AP00004.

## Materials and Methods

### Ethics statement

All mouse studies followed the Guide for the Care and Use of Laboratory of Animals of the National Institutes of Health. The animal protocols (2015-0315 D and 2018-0488 D) were reviewed and approved by Texas A&M Institutional Animal Care and Use Committee (IACUC).

### Bacterial strains and media

Methicillin-resistant *Staphylococcus aureus* isolate used in this study was the kind gift of Dr. Magnus Hook (Texas A&M Institute of Biosciences and Technology, Houston). USA300 is a fully virulent, community-acquired clone of MRSA. Strains were routinely cultured in Luria-Bertani (LB) broth and plates supplemented with antibiotics when needed at 50 mg/L Kanamycin Sulphate. For murine infections, strains were grown aerobically at 37^0^C to a stationary phase in LB broth supplemented with Kanamycin.

### Murine strains

Both conventional (C57BL/6J) and Collaborative Cross mice were used for these experiments. In total, three males and three females belonging to 25 different CC strains (chosen at random) and C57BL/6J were used (156 mice total). All mice were initially purchased from UNC’s Systems Genetics Core Facility (SGCF). They were bred independently at the Division of Comparative Medicine at Texas A&M University (Table S1). Mice were fed Envigo Teklad Global 19% Protein Extrudent Rodent Diet (2919) or Envigo Teklad Rodent Diet (8604) based on strain need. Mice were provided with cardboard huts, food, and water ad libitum.

### Placement of telemetry devices

Mice (5-7 weeks old) were anesthetized with an isoflurane vaporizer using the Kent Scientific SomnoSuite Low-flow anesthesia system to implant telemetry devices. A midline abdominal incision was made, and Starr Life Science G2 Emitter devices were sutured to the ventral abdominal wall. The abdominal muscle layer was sutured with 5-0 vicryl, and the skin layer was closed using stainless steel wound clips. Buprenorphine (0.0001 mg/g) was administered intraperitoneally before recovery from anesthesia. Mice were monitored twice daily for surgical complications and humanely euthanized when needed. Surgical clips were removed on the seventh day, and animals were allowed to recover for one more week.

### Infection with MRSA

After baseline scoring for health, 6 mice (now 8-12 weeks old) per CC strain were infected with MRSA USA300. We chose a 7-day screening period and very sensitive telemetry monitoring of MRSA infected animals in an attempt to capture and identify range of disease phenotypes from highly susceptible to resistant. To minimize batch effects, mice from a given CC strain were infected in different experiments across a two-year period. This approach may have resulted in some within strain variability in infection outcomes. Briefly, Mice were anesthetized using isoflurane and infected with 1 x 10^7^ CFU in 50 µl of LB broth intravenously at the inferior fornix into the retro-orbital sinus. For each experiment, mice were inoculated at the same time of day. Mice that became moribund within 6 hours of infection were humanely euthanized and removed from the experiment.

### Health monitoring

After the recovery from surgery, mice were moved to a BSL-2 facility and acclimated for 5-7 days. Individual cages containing implanted mice were placed onto ER4000 receiver platforms and calibrated to receive signals from the implanted telemetry devices. The outputs, body temperature (once per minute) and gross motor activity (summation of movement per minute), were continuously fed to a computer system for visualization (Fig S5 and Fig S6). This sensitive continuous monitoring allowed us understand disease progression for each animal in real-time, as disruptions of the normal circadian pattern of core body temperature after infection indicated symptomatic MRSA infection. A machine-learning algorithm was used to identify the time to deviation from the circadian pattern of body temperature and activity. Detailed explanation of the methodology and calculations can be found here [51].

We manually scored four health parameters twice daily: physical appearance, body conditioning, and provoked and unprovoked behavior. The scoring scale ranged from 0-3, with zero = normal and three = abnormal (Table S2). Additionally, as a measure of activity, four small nestlets were placed at each corner of the cage in the evening. The following day, the number of nestlets moved from the cage corners into the hut was noted.

### Euthanasia criteria

After infection, mice were monitored continuously using telemetry readings and twice daily using visual health scoring. For telemetry, euthanasia criteria were defined by a sudden drop in temperature (3^0^C or more). For manual heath scoring, the euthanasia criteria were reached when the combined health score exceeded 8. Mice that met one or both these criteria were humanely euthanized by CO2 asphyxiation.

### Bacterial load determination

After euthanasia, whole blood, serum, and organs (spleen, liver, heart, lung, and kidney) were collected. A consistent region of each organ was collected in 3mL ice-cold PBS and homogenized. The serially diluted homogenate was plated on LB plates supplemented with kanamycin for bacterial enumeration in each organ. Data are expressed as CFU/g of tissue. Total organ colonization was calculated by adding CFU/g values of all the five organs collected.

### Histopathology

After euthanasia, portions of collected organs were fixed in 10% neutral buffered formalin for 24 hours and stored in 70% ethanol. Fixed tissue was embedded in paraffin, sectioned at 5 µm, and stained with hematoxylin and eosin (H&E). A board-certified veterinary pathologist scored all the slides for tissue damage on a scale of 0 to 4 in a blinded manner (Table S3). All the raw pathology data is available in the supplementary file (Table S2). Whole slide images of H&E-stained tissue sections were captured as digital files by scanning at 40X using a 3D Histech Pannoramic SCAN II FL Scanner (Epredia, MI). Digital files were processed by Aiforia Hub (Cambridge, MA) software to generate images with scaled bars.

### Complete blood count

One week before the implantation of telemetry devices, whole blood was collected from each mouse by submandibular bleeding. From the same mice, blood was collected at necropsy after infection by cardiac puncture. Abaxis VetScan HM5, optimized for rodents, was used to analyze blood collected in Ethylene Diamine Tetra Acetic acid (EDTA) tubes. The ratio between the blood parameters after and before infection (value >1 = increased after infection and value < 1 = decreased after infection) was calculated and reported.

### Heritability

Broad-sense heritability (H^2^) was calculated using the formula H^2^ = V_G_/V_P_ = V_G_/(V_E_ + V_G_) as previous described [114, 115]. V_E_ and V_G_ are the variance explained by the environmental and the genetic component respectively while V_P_ is the total phenotypic variance for a given phenotype. Detailed explanation of the calculation can be found here [116].

### QTL analysis

QTL analysis was performed using R/qtl2 software [59]. This method accounts for the complex population structure in CC strains. Briefly, genotype probabilities were imputed from the QTL viewer [60]. Genome scans were performed on the transformed phenotype using the scan1 function. The generated Logarithm of Odds (LOD) score is the likelihood ratio comparing the hypothesis of a QTL at a given position versus that of no QTL. We used the number of mice within a strain that survived at the end of each day as the phenotype. The phenotype was randomly shuffled 999 times to establish genome-wide significance, and LOD scores were calculated for each iteration using the scan1perm function [117]. The 85^th^ percentile of the scores was considered significant for that phenotype. The genomic confidence interval was calculated by dropping the LOD scores by 1.8 for each significant peak. Mouse Genome Informatics (MGI) was used to find the genes and QTL features within each interval using mouse genome version GRCm38 [118]. To further shortlist candidate genes, the founder strain distribution pattern was queried against the CC variant database (V3 Version). The Variant effect predictor [VEP] from the ensemble genome database was used to calculate the impact score for the variants [119].

### RNA extraction and sequencing

Total RNA was extracted from frozen tissues using Direct-zol RNA Miniprep plus kit following the manufacturer’s protocol (Zymo Research - R2073). The purity and quantity of the extracted RNA were analyzed using RNA 6000 Nano LabChip Kit and Bioanalyzer 2100 (Agilent CA, USA, 5067-1511). High-Quality RNA samples with RIN number > 7.0 were used to construct the sequencing library. After total RNA was extracted, mRNA was purified from total RNA (5µg) using Dynabeads Oligo (dT) with two rounds of purification (Thermo Fisher, CA, USA). Following purification, the mRNA was fragmented into short fragments using divalent cations under elevated temperature (Magnesium RNA Fragmentation Module (NEB, cat. e6150, USA) under 94℃ 5-7min). Then the cleaved RNA fragments were reverse-transcribed to create the cDNA by SuperScript™ II Reverse Transcriptase (Invitrogen, cat. 1896649, USA), which were next used to synthesize U-labeled second-stranded DNAs with E. coli DNA polymerase I (NEB, cat.m0209, USA), RNase H (NEB, cat.m0297, USA) and dUTP Solution (Thermo Fisher, cat. R0133, USA). An A-base was then added to the blunt ends of each strand, preparing them for ligation to the indexed adapters. Each adapter contained a T-base overhang for ligating the adapter to the A-tailed fragmented DNA. Dual-index adapters were ligated to the fragments, and size selection was performed with AMPureXP beads. After the heat-labile UDG enzyme (NEB, cat.m0280, USA) treatment of the U-labeled second-stranded DNAs, the ligated products were amplified with PCR by the following conditions: initial denaturation at 95℃ for 3 min; 8 cycles of denaturation at 98℃ for 15 sec, annealing at 60℃ for 15 sec, and extension at 72℃ for 30 sec; and then final extension at 72℃ for 5 min. The average insert size for the final cDNA libraries was 300±50 bp. At last, we performed the 2×150bp paired-end sequencing (PE150) on an Illumina Novaseq™ 6000 following the vendor’s recommended protocol.

### RNA sequencing data analysis

Reads were trimmed using trim galore (version 0.6.7) [120–123]. This removed adapters, poly tails, more than 5% of unknown nucleotides, and low-quality reads containing more than 20% of low-quality bases (Q-value <20). Both forward and reverse hard trimmed at 100 base pairs. FastQC was used to verify data quality before and after cleaning [123]. Cleaned reads were aligned and counted against the mouse reference genome (GRCm39) using STAR (version 2.7.9a) aligner [124]. Downstream processing of the data was performed using IDEP 1.0 [125, 126]. Gene counts were analyzed for differentially expressed genes using DESeq2 [127]

### Data availability

All of the data points collected on each mouse before and after the infection can be found in the supplementary file (Table S2). Raw RNA sequencing files can be found at SRA (PRJNA1002380).

