## Supplementary material for "Collaborative Cross mice have diverse phenotypic responses to infection with Methicillin-resistant *Staphylococcus aureus* USA300": Fig S1-S4

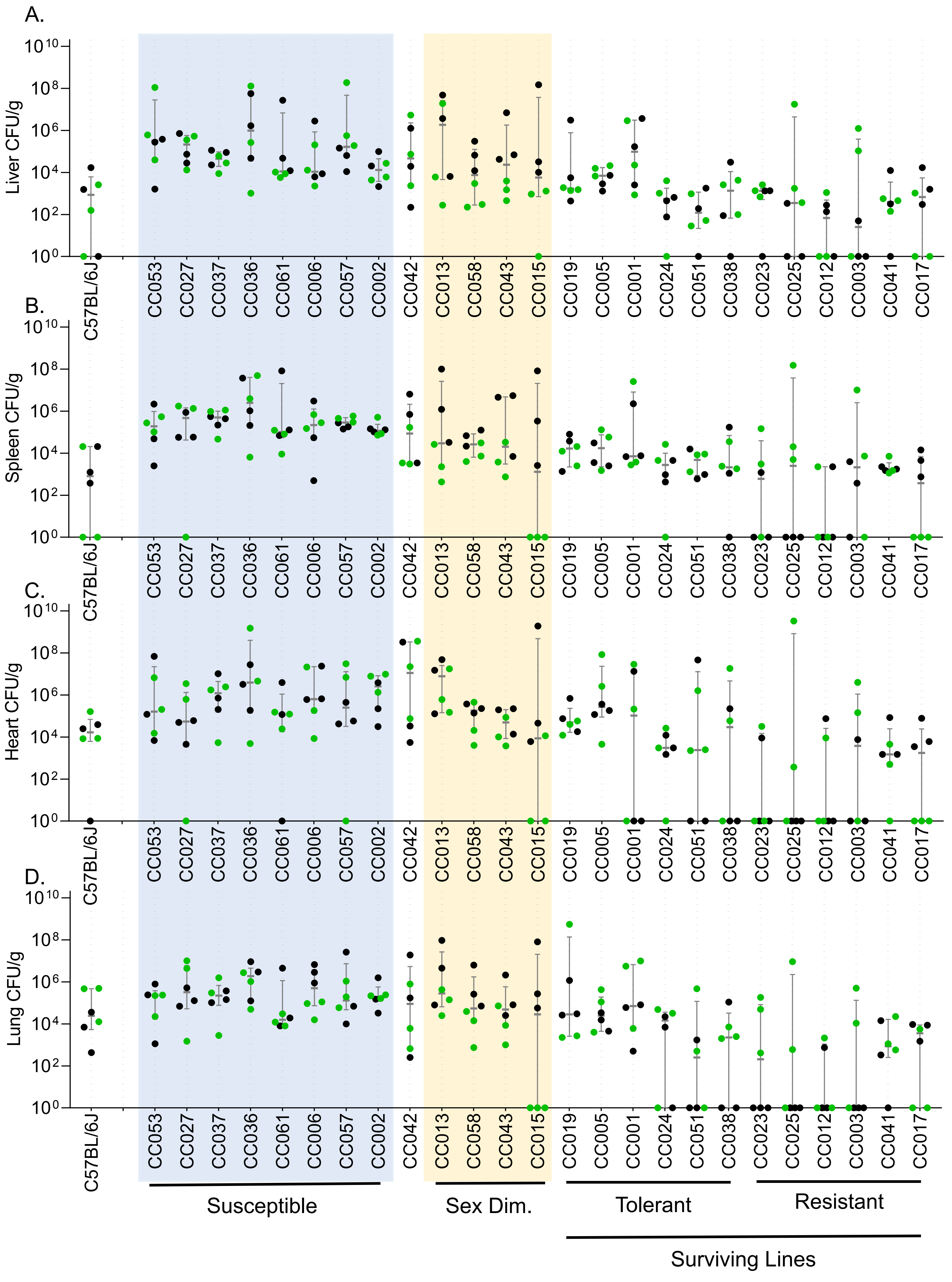
A.
B.
C.
D.
Liver CFU/g
Spleen CFU/g
Heart CFU/g
Lung CFU/g
$10^{0}$
$10^{2}$
$10^{4}$
$10^{6}$
$10^{8}$
$10^{10}$
C57BL/6J
CC053
CC027
CC037
CC036
CC061
CC006
CC057
CC002
CC042
CC013
CC058
CC043
CC015
CC019
CC005
CC001
CC024
CC051
CC038
CC023
CC025
CC012
CC003
CC041
CC017
Susceptible
Sex Dim.
Tolerant
Resistant
Surviving Lines

Fig S2

Spleen Ordinal Scoring

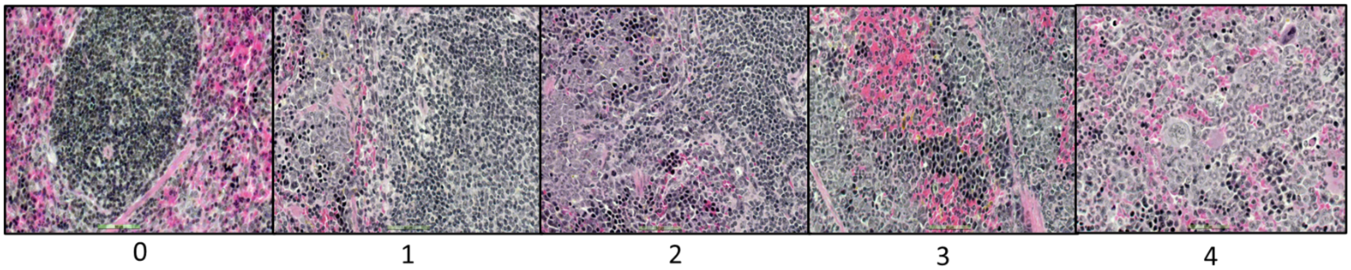


Liver Ordinal Scoring

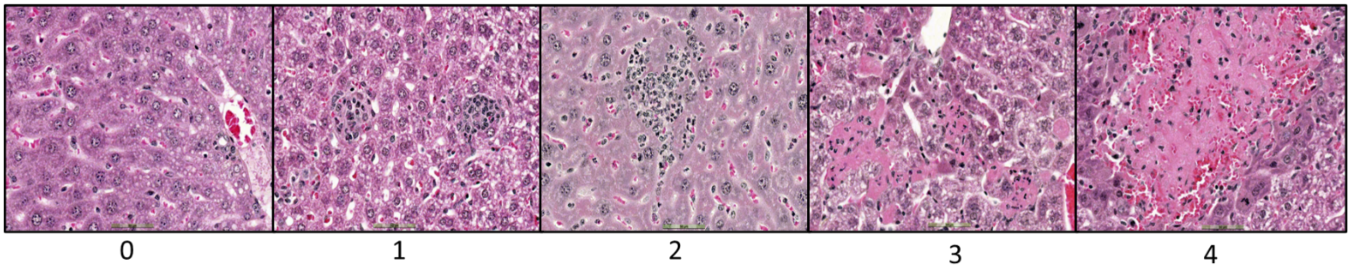


Heart Ordinal Scoring

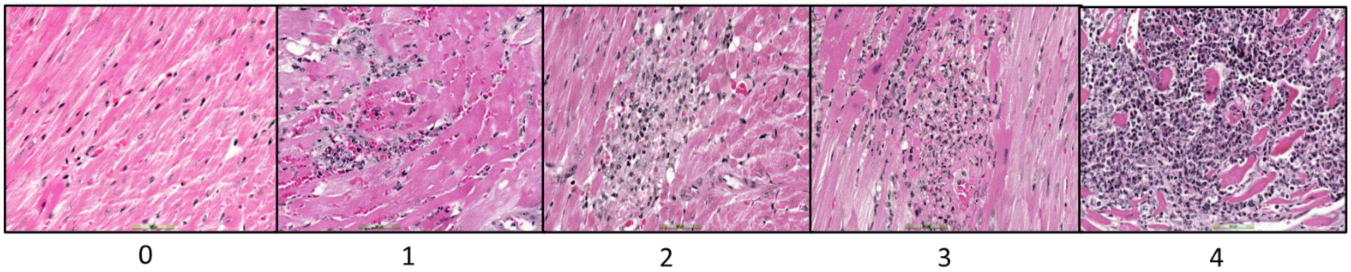


Lung Ordinal Scoring

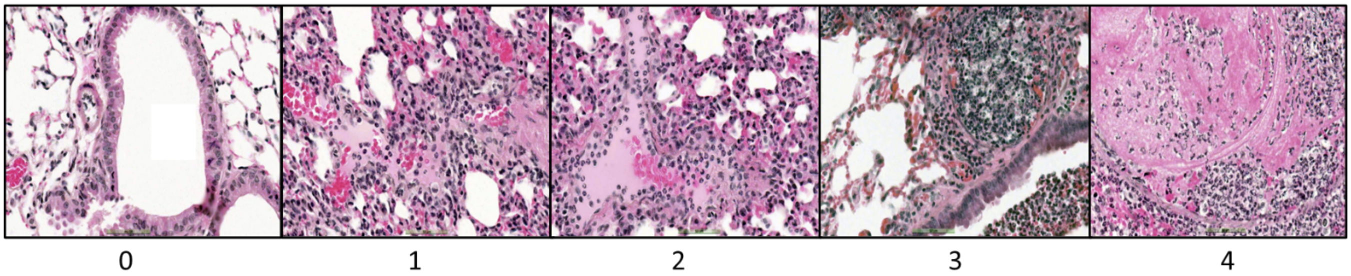


Kidney Ordinal Scoring

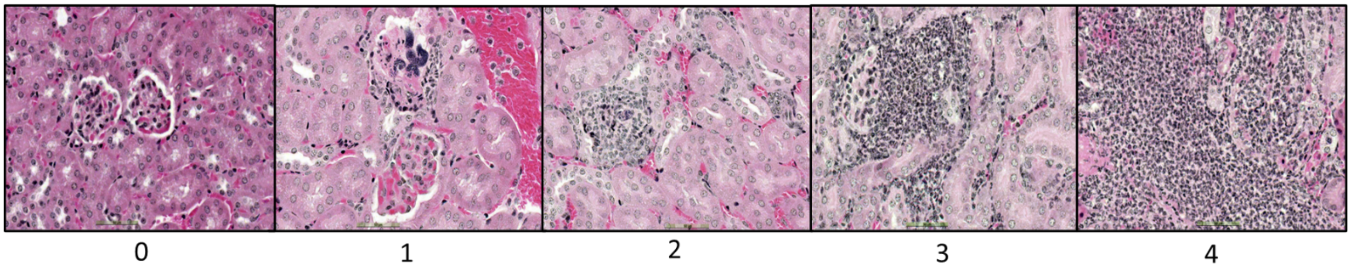

Fig S3

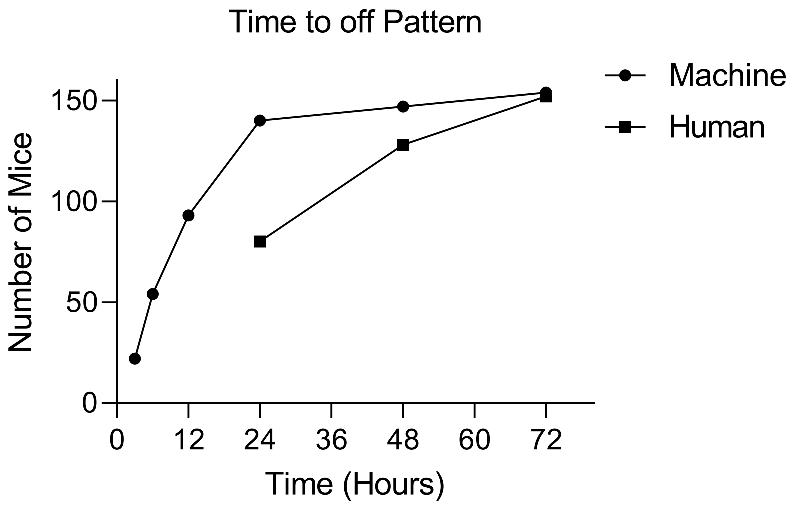

Time to off Pattern
Machine
Human
Number of Mice
150
100
50
0
0
12
24
36
48
60
72
Time (Hours)

Fig S4

A.

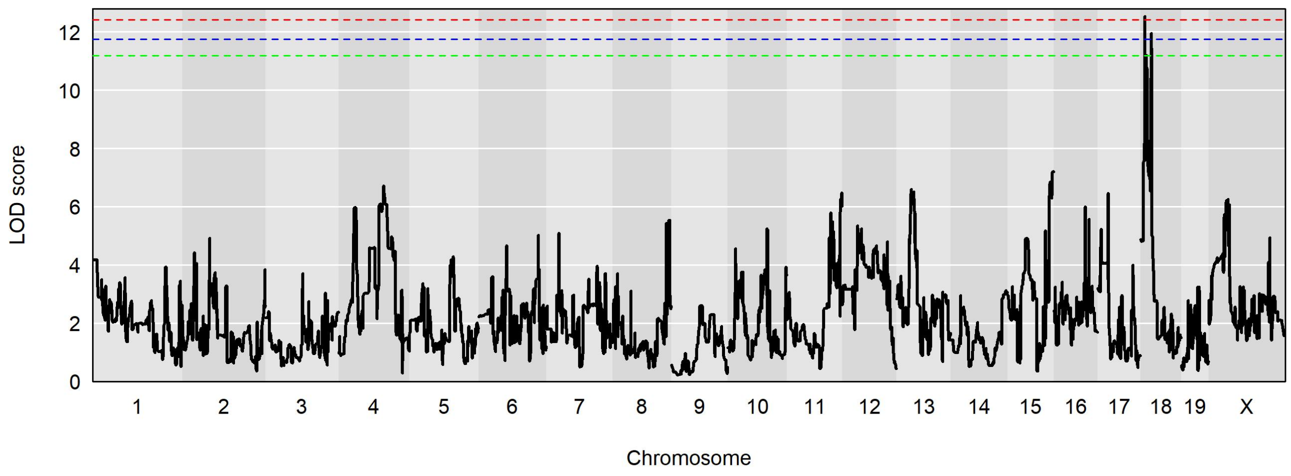
LOD score
0
2
4
6
8
10
12
1 2 3 4 5 6 7 8 9 10 11 12 13 14 15 16 17 18 19 X
Chromosome

B.

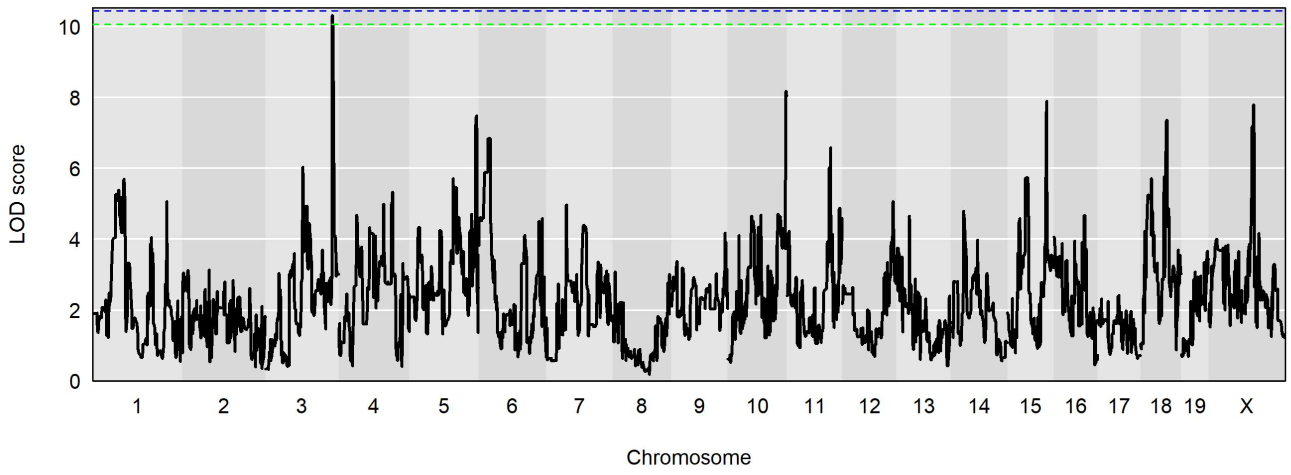
LOD score
0
2
4
6
8
10
1 2 3 4 5 6 7 8 9 10 11 12 13 14 15 16 17 18 19 X
Chromosome
