## Supplementary material for "Collaborative Cross mice have diverse phenotypic responses to infection with Methicillin-resistant *Staphylococcus aureus* USA300": Fig S5

Supplementary Fig S8

Circadian pattern for all mice involved in the study. Blue line represents temperature, black line represents activity and red line represents the time of infection.

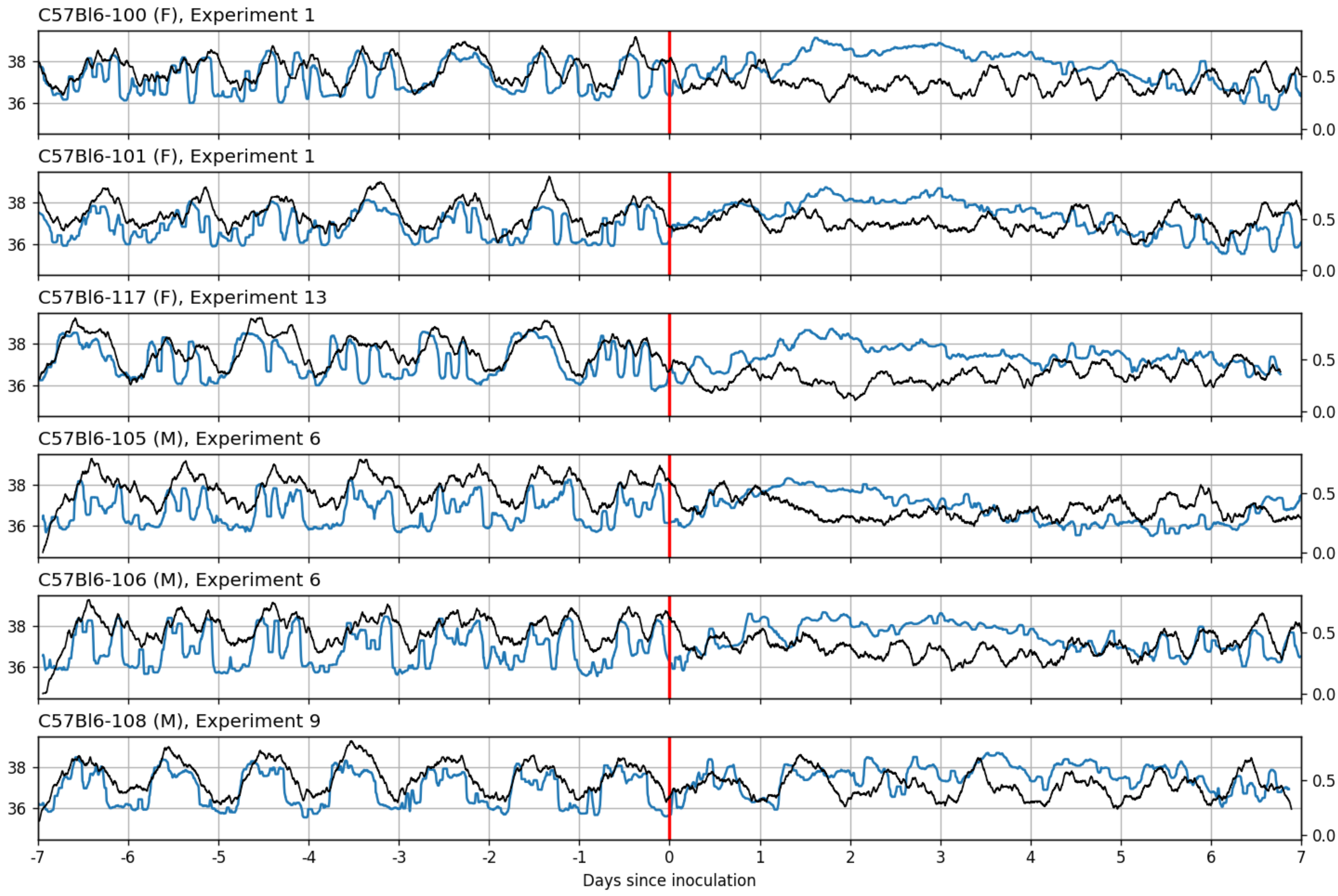

C57Bl6-100 (F), Experiment 1
C57Bl6-101 (F), Experiment 1
C57Bl6-117 (F), Experiment 13
C57Bl6-105 (M), Experiment 6
C57Bl6-106 (M), Experiment 6
C57Bl6-108 (M), Experiment 9
38
36
0.5
0.0
-7
-6
-5
-4
-3
-2
-1
0
1
2
3
4
5
6
7
Days since inoculation

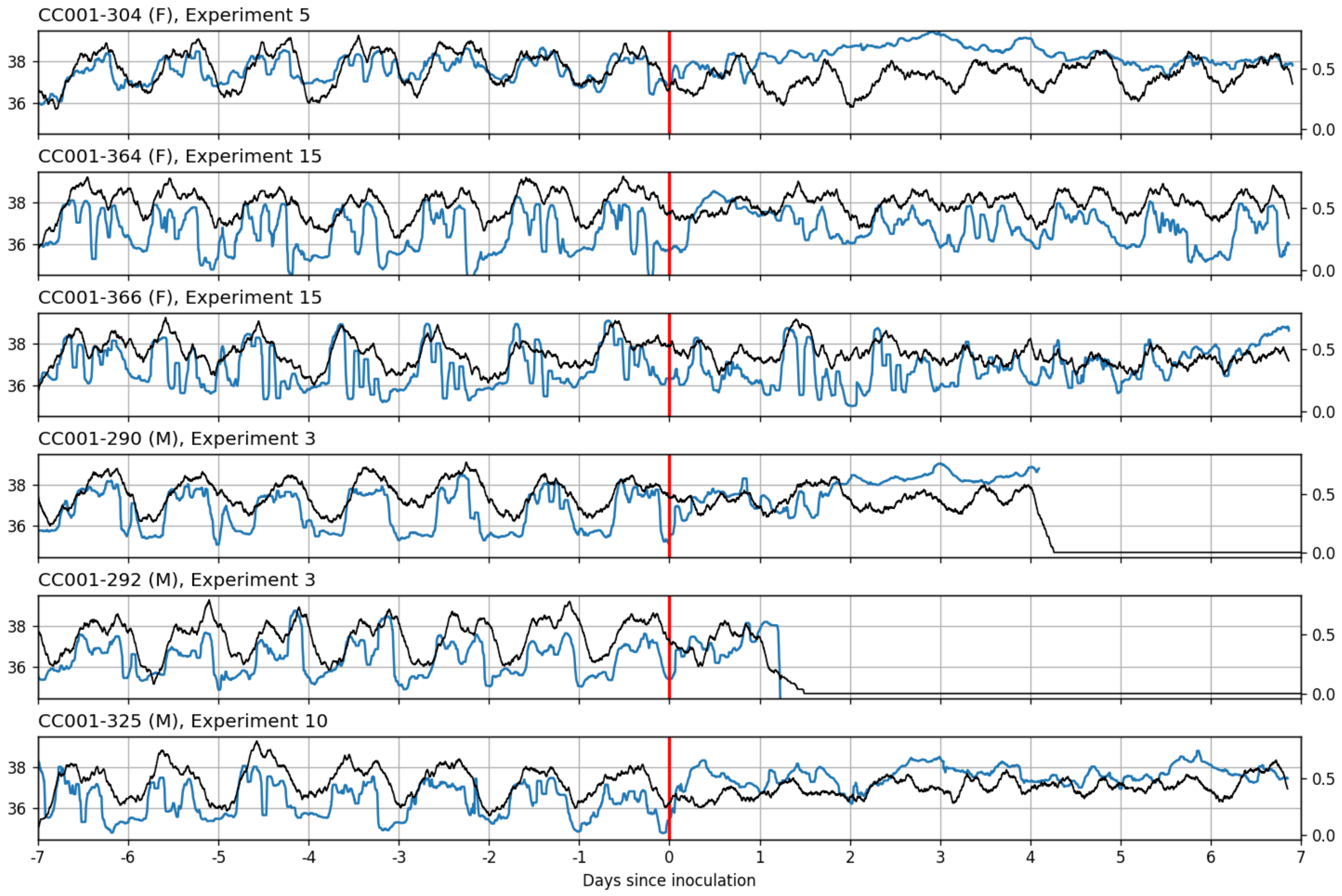

CC001-304 (F), Experiment 5
CC001-364 (F), Experiment 15
CC001-366 (F), Experiment 15
CC001-290 (M), Experiment 3
CC001-292 (M), Experiment 3
CC001-325 (M), Experiment 10
38
36
0.5
0.0
-7
-6
-5
-4
-3
-2
-1
0
1
2
3
4
5
6
7
Days since inoculation

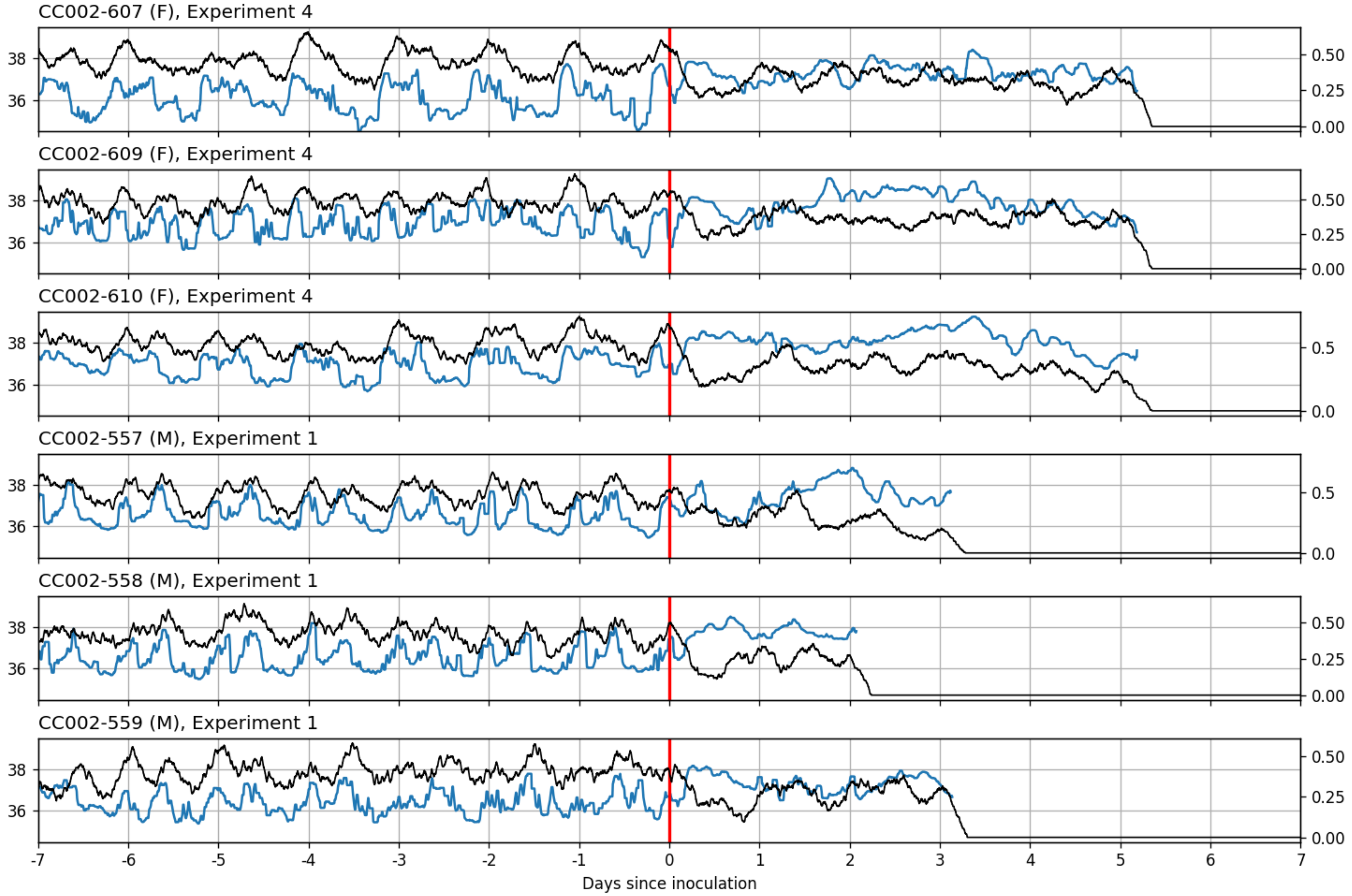

CC002-607 (F), Experiment 4
CC002-609 (F), Experiment 4
CC002-610 (F), Experiment 4
CC002-557 (M), Experiment 1
CC002-558 (M), Experiment 1
CC002-559 (M), Experiment 1
Days since inoculation

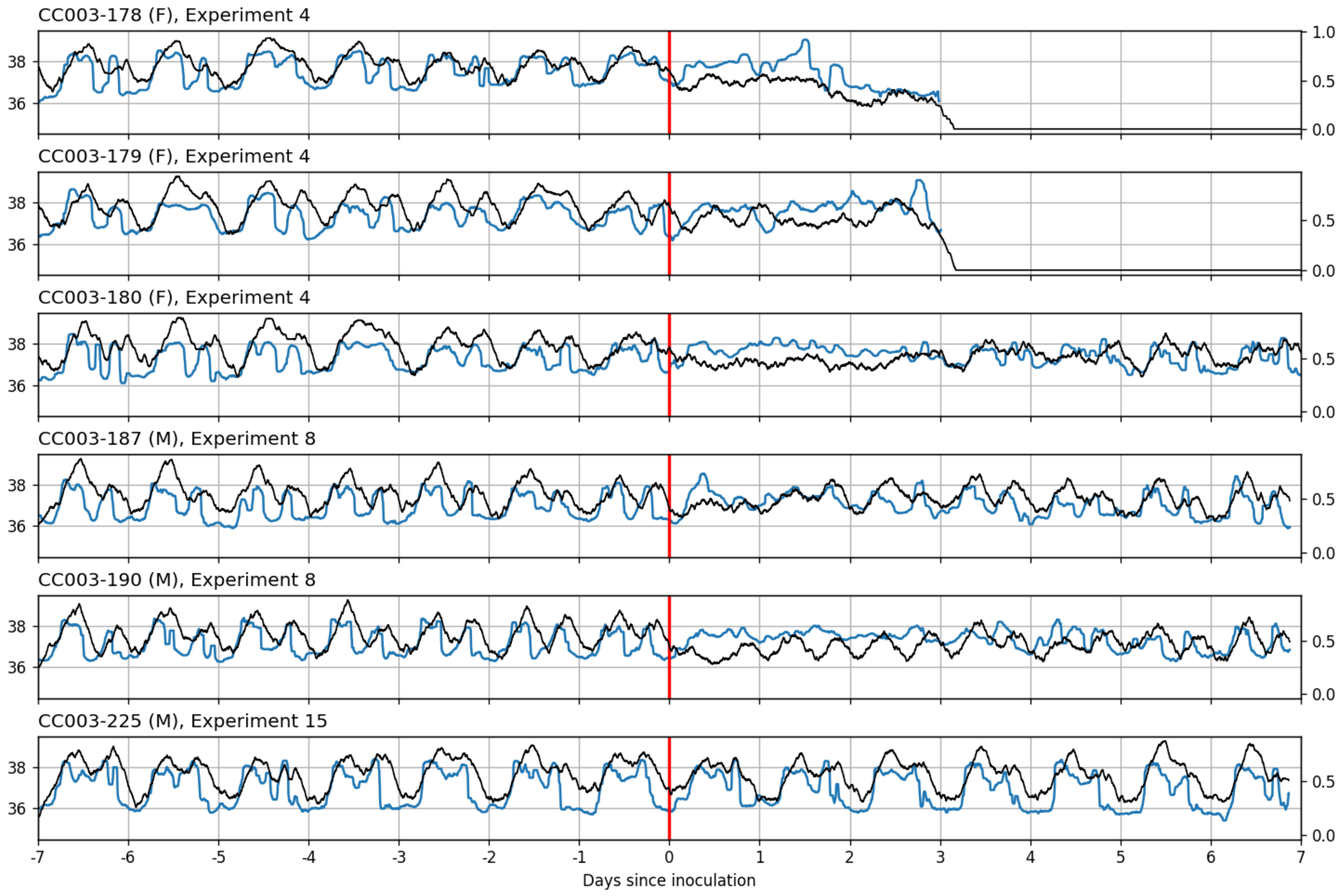

CC003-178 (F), Experiment 4
CC003-179 (F), Experiment 4
CC003-180 (F), Experiment 4
CC003-187 (M), Experiment 8
CC003-190 (M), Experiment 8
CC003-225 (M), Experiment 15
36
38
0.0
0.5
1.0
-7
-6
-5
-4
-3
-2
-1
0
1
2
3
4
5
6
7
Days since inoculation

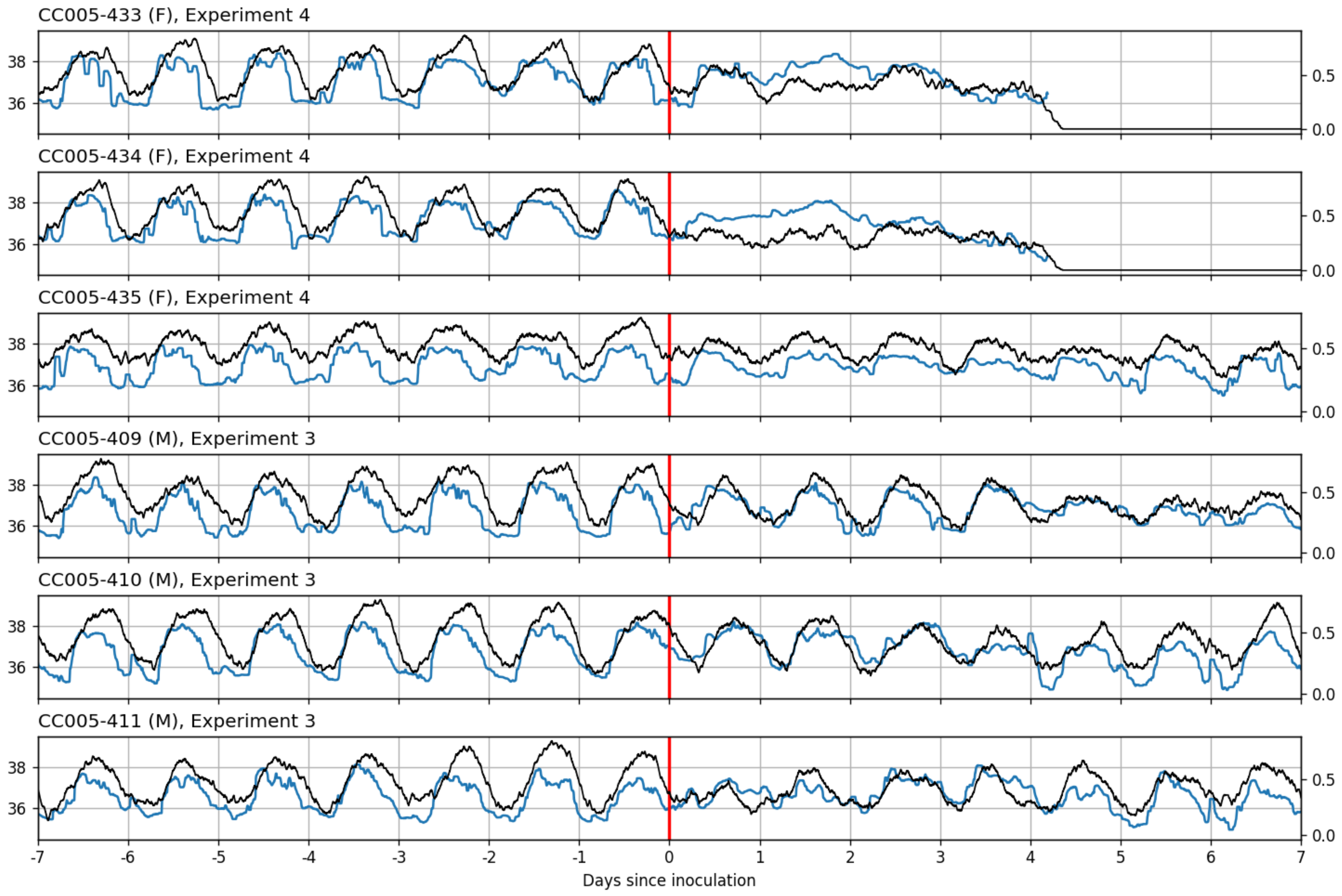

CC005-433 (F), Experiment 4
CC005-434 (F), Experiment 4
CC005-435 (F), Experiment 4
CC005-409 (M), Experiment 3
CC005-410 (M), Experiment 3
CC005-411 (M), Experiment 3
36
38
0.0
0.5
-7
-6
-5
-4
-3
-2
-1
0
1
2
3
4
5
6
7
Days since inoculation

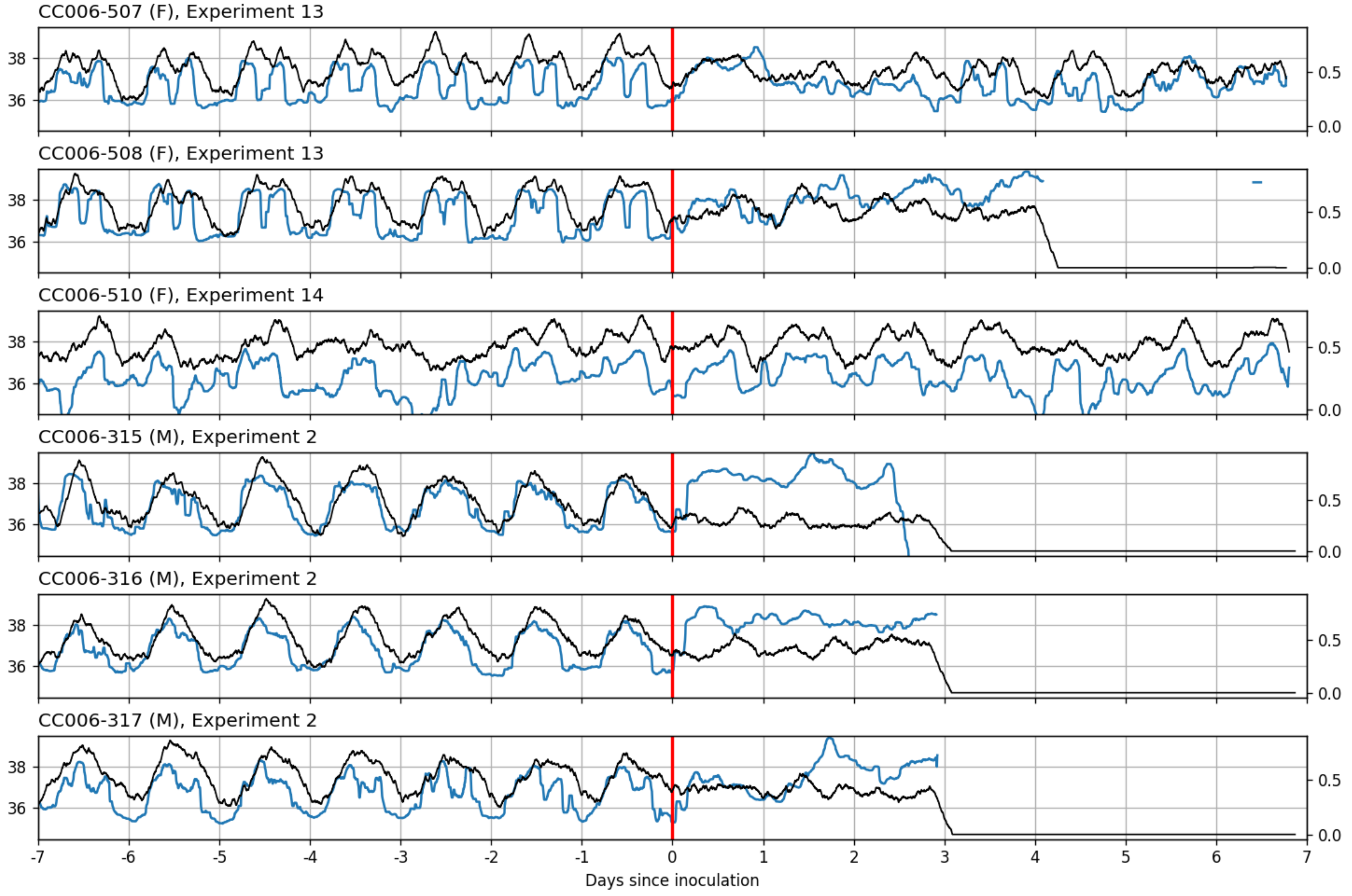

CC006-507 (F), Experiment 13
CC006-508 (F), Experiment 13
CC006-510 (F), Experiment 14
CC006-315 (M), Experiment 2
CC006-316 (M), Experiment 2
CC006-317 (M), Experiment 2
38
36
0.5
0.0
-7
-6
-5
-4
-3
-2
-1
0
1
2
3
4
5
6
7
Days since inoculation

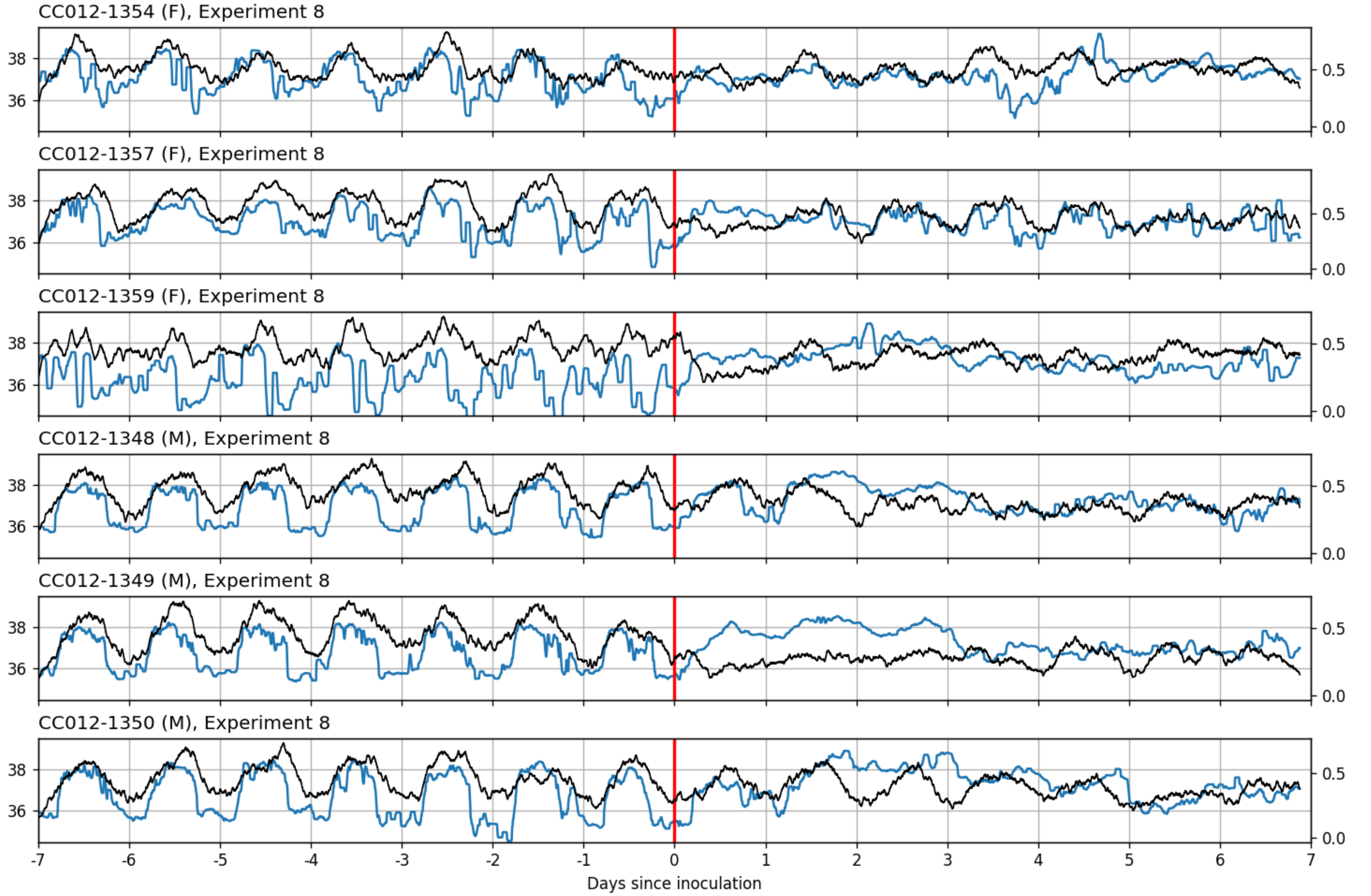
CC012-1354 (F), Experiment 8
CC012-1357 (F), Experiment 8
CC012-1359 (F), Experiment 8
CC012-1348 (M), Experiment 8
CC012-1349 (M), Experiment 8
CC012-1350 (M), Experiment 8
36
38
0.0
0.5
-7
-6
-5
-4
-3
-2
-1
0
1
2
3
4
5
6
7
Days since inoculation

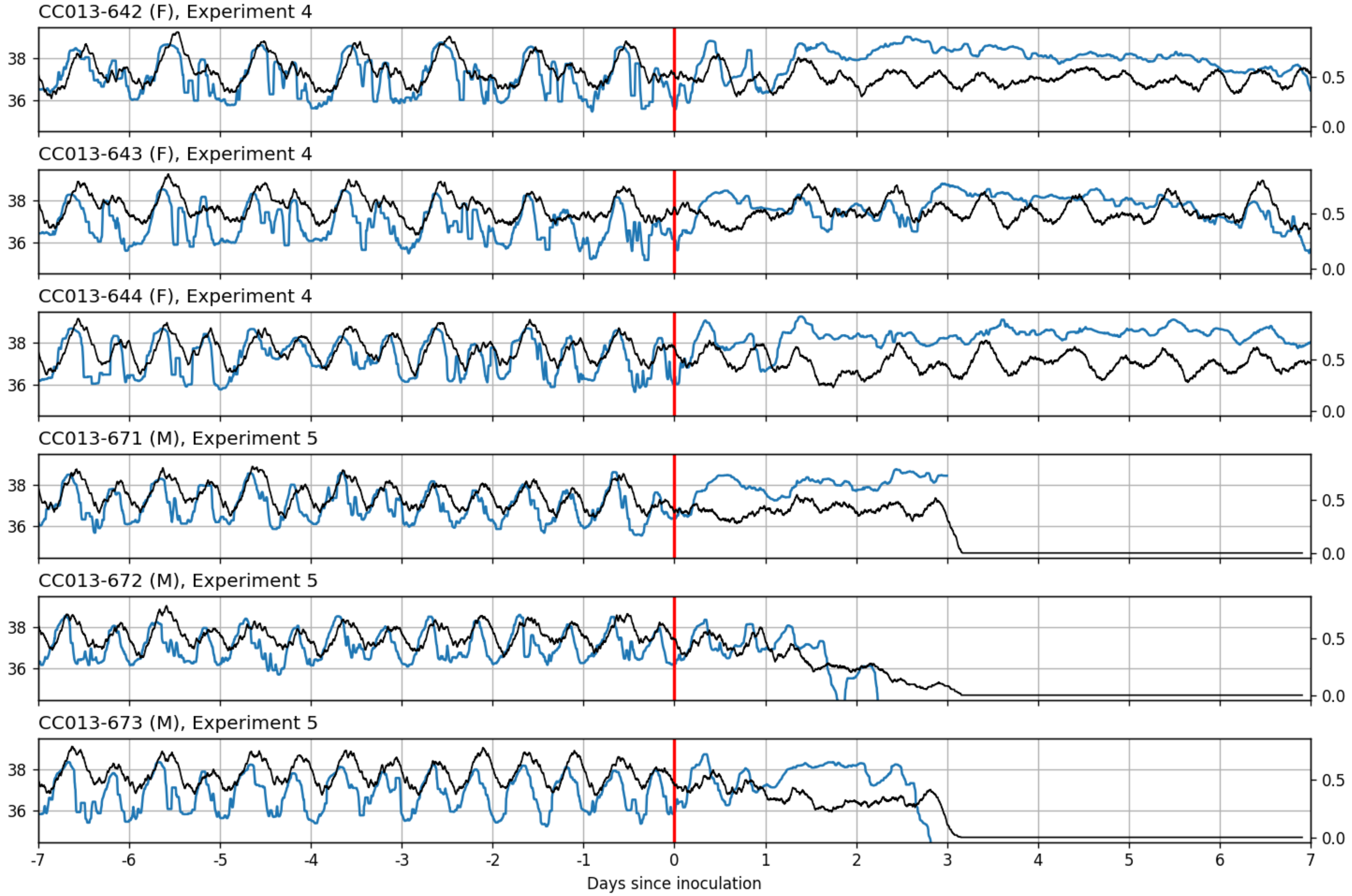
CC013-642 (F), Experiment 4
CC013-643 (F), Experiment 4
CC013-644 (F), Experiment 4
CC013-671 (M), Experiment 5
CC013-672 (M), Experiment 5
CC013-673 (M), Experiment 5
38
36
0.5
0.0
-7
-6
-5
-4
-3
-2
-1
0
1
2
3
4
5
6
7
Days since inoculation

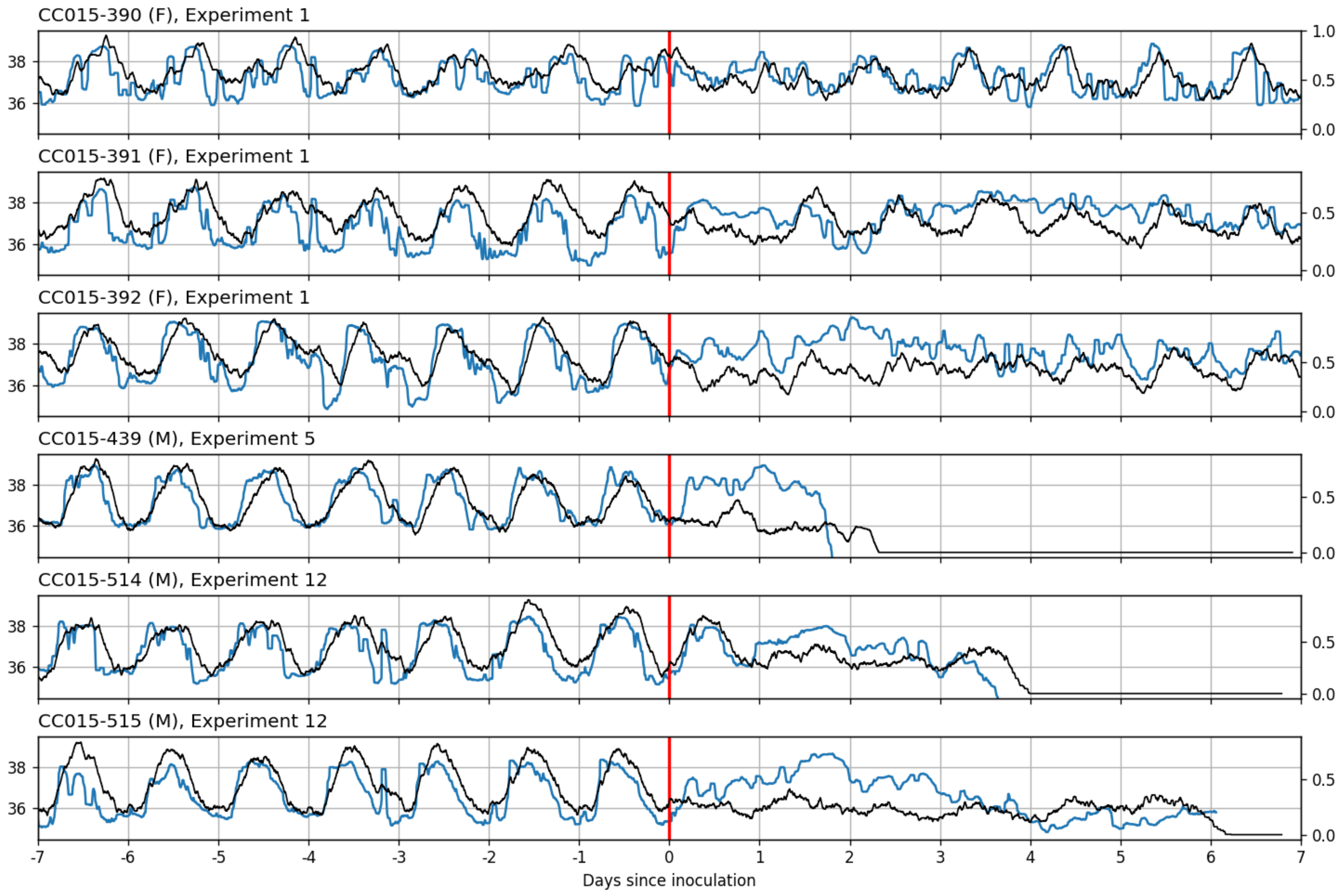

CC015-390 (F), Experiment 1
CC015-391 (F), Experiment 1
CC015-392 (F), Experiment 1
CC015-439 (M), Experiment 5
CC015-514 (M), Experiment 12
CC015-515 (M), Experiment 12
38
36
1.0
0.5
0.0
-7
-6
-5
-4
-3
-2
-1
0
1
2
3
4
5
6
7
Days since inoculation

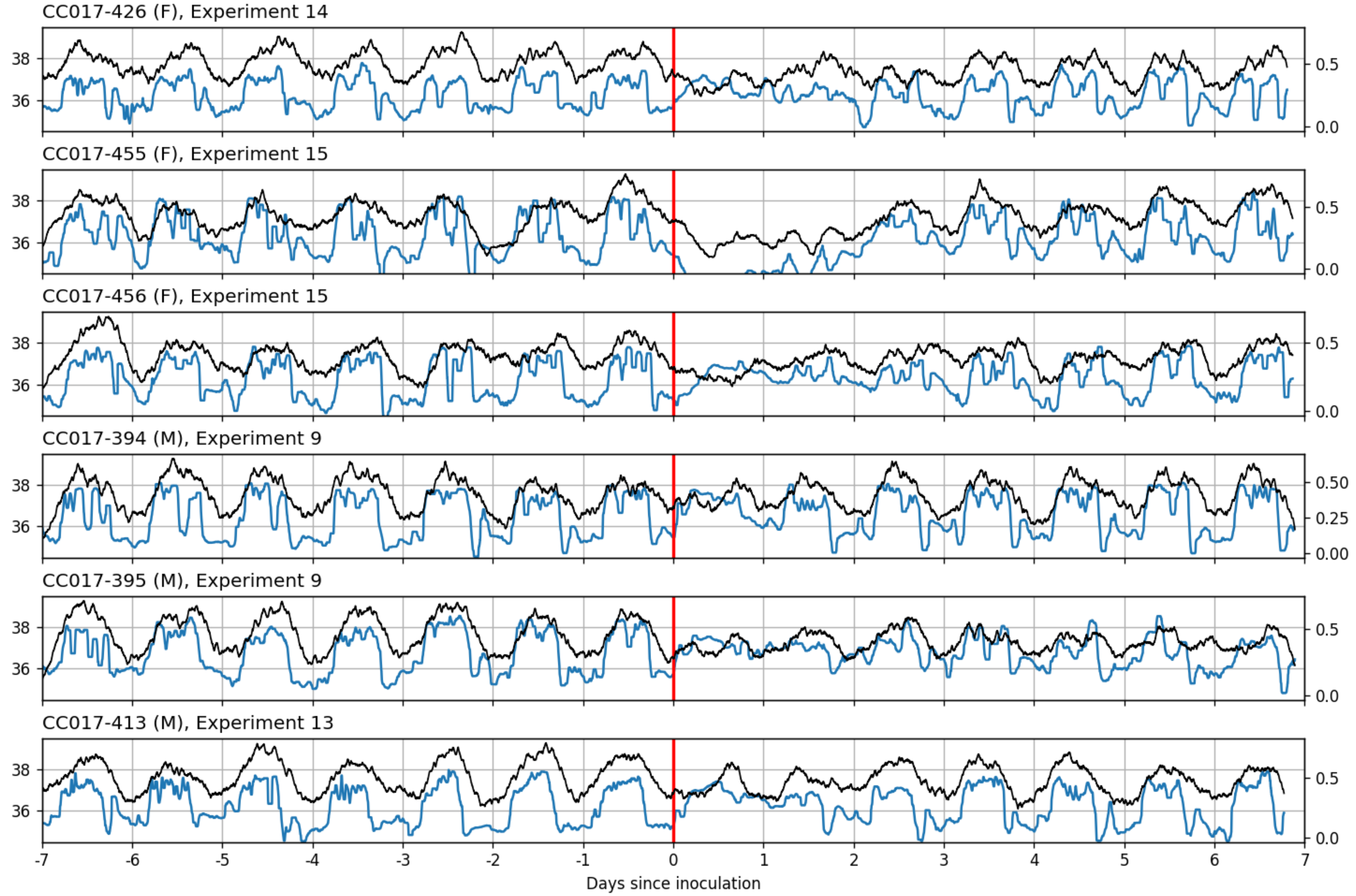

CC017-426 (F), Experiment 14
CC017-455 (F), Experiment 15
CC017-456 (F), Experiment 15
CC017-394 (M), Experiment 9
CC017-395 (M), Experiment 9
CC017-413 (M), Experiment 13
Days since inoculation

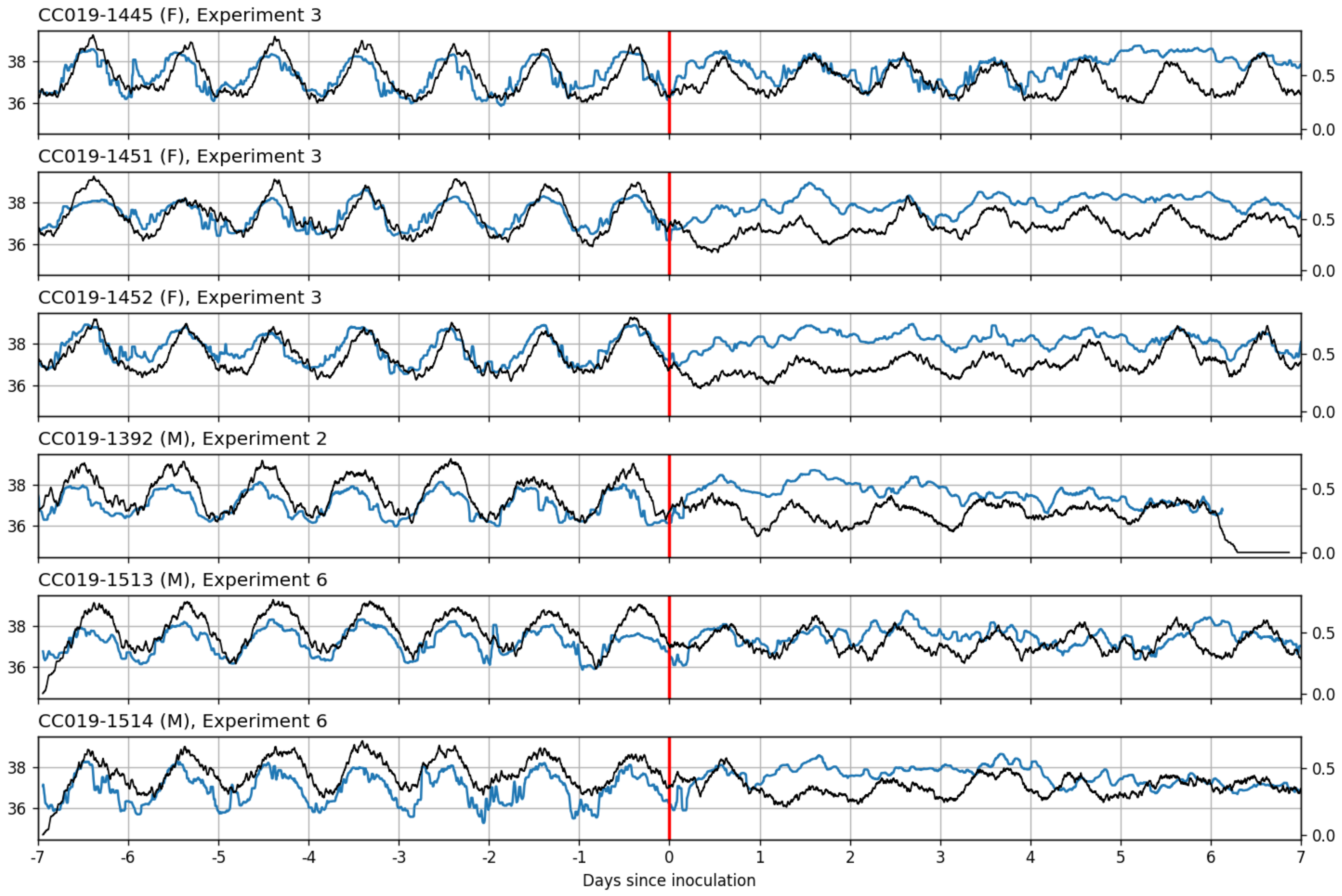
CC019-1445 (F), Experiment 3
CC019-1451 (F), Experiment 3
CC019-1452 (F), Experiment 3
CC019-1392 (M), Experiment 2
CC019-1513 (M), Experiment 6
CC019-1514 (M), Experiment 6
36
38
0.0
0.5
-7
-6
-5
-4
-3
-2
-1
0
1
2
3
4
5
6
7
Days since inoculation

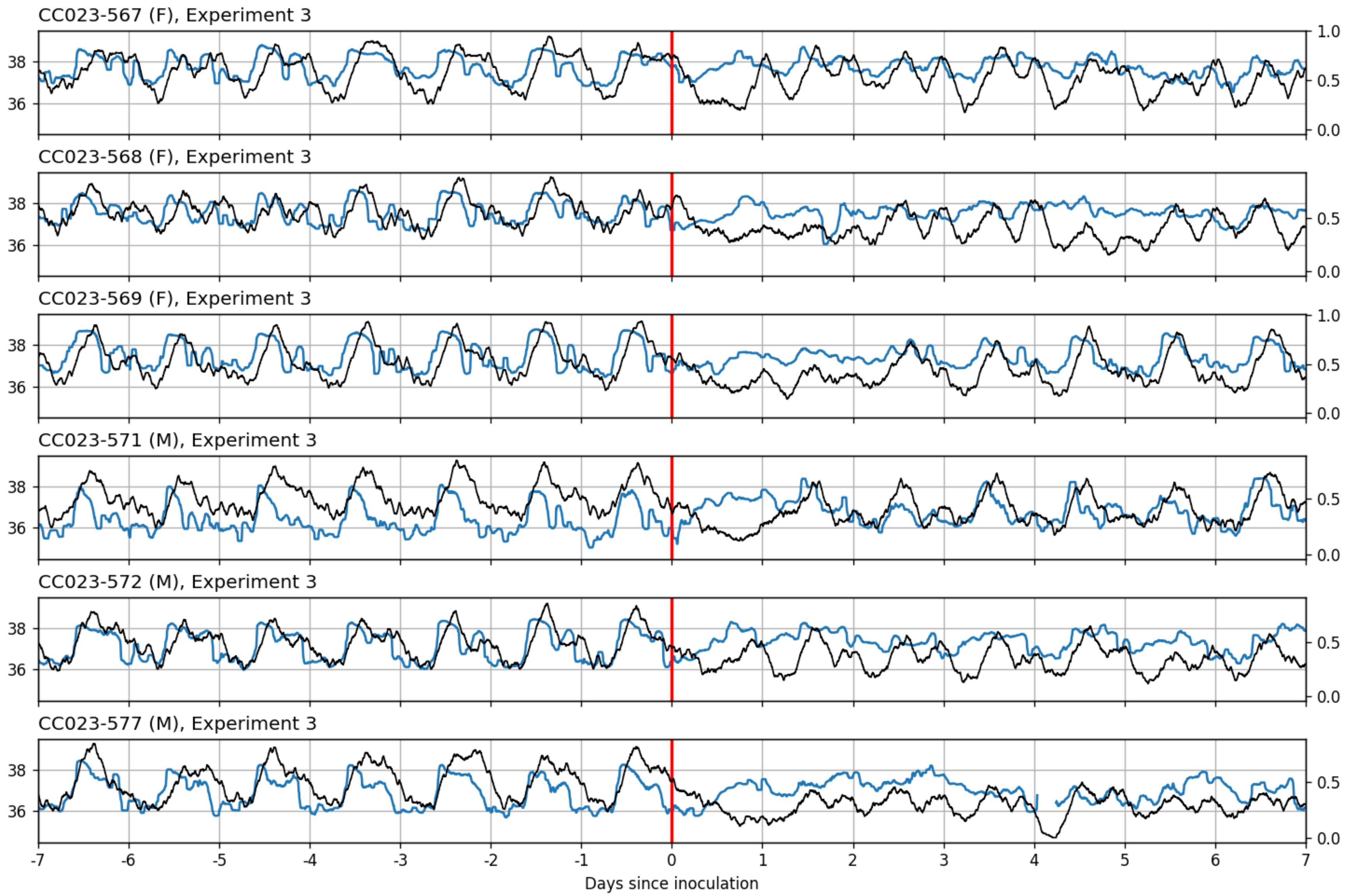
CC023-567 (F), Experiment 3
CC023-568 (F), Experiment 3
CC023-569 (F), Experiment 3
CC023-571 (M), Experiment 3
CC023-572 (M), Experiment 3
CC023-577 (M), Experiment 3
38
36
1.0
0.5
0.0
-7
-6
-5
-4
-3
-2
-1
0
1
2
3
4
5
6
7
Days since inoculation

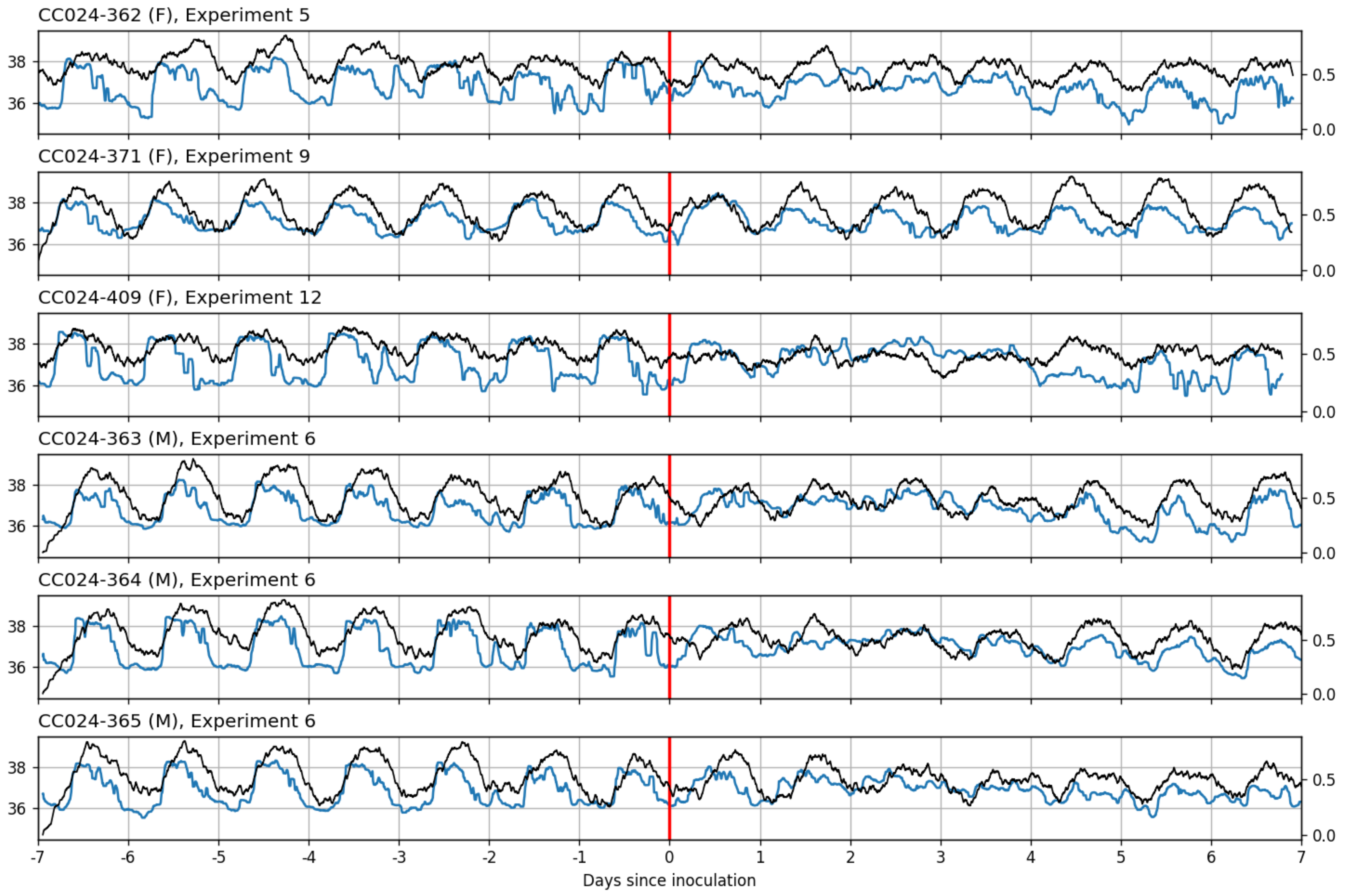
CC024-362 (F), Experiment 5
CC024-371 (F), Experiment 9
CC024-409 (F), Experiment 12
CC024-363 (M), Experiment 6
CC024-364 (M), Experiment 6
CC024-365 (M), Experiment 6
36
38
0.0
0.5
-7
-6
-5
-4
-3
-2
-1
0
1
2
3
4
5
6
7
Days since inoculation

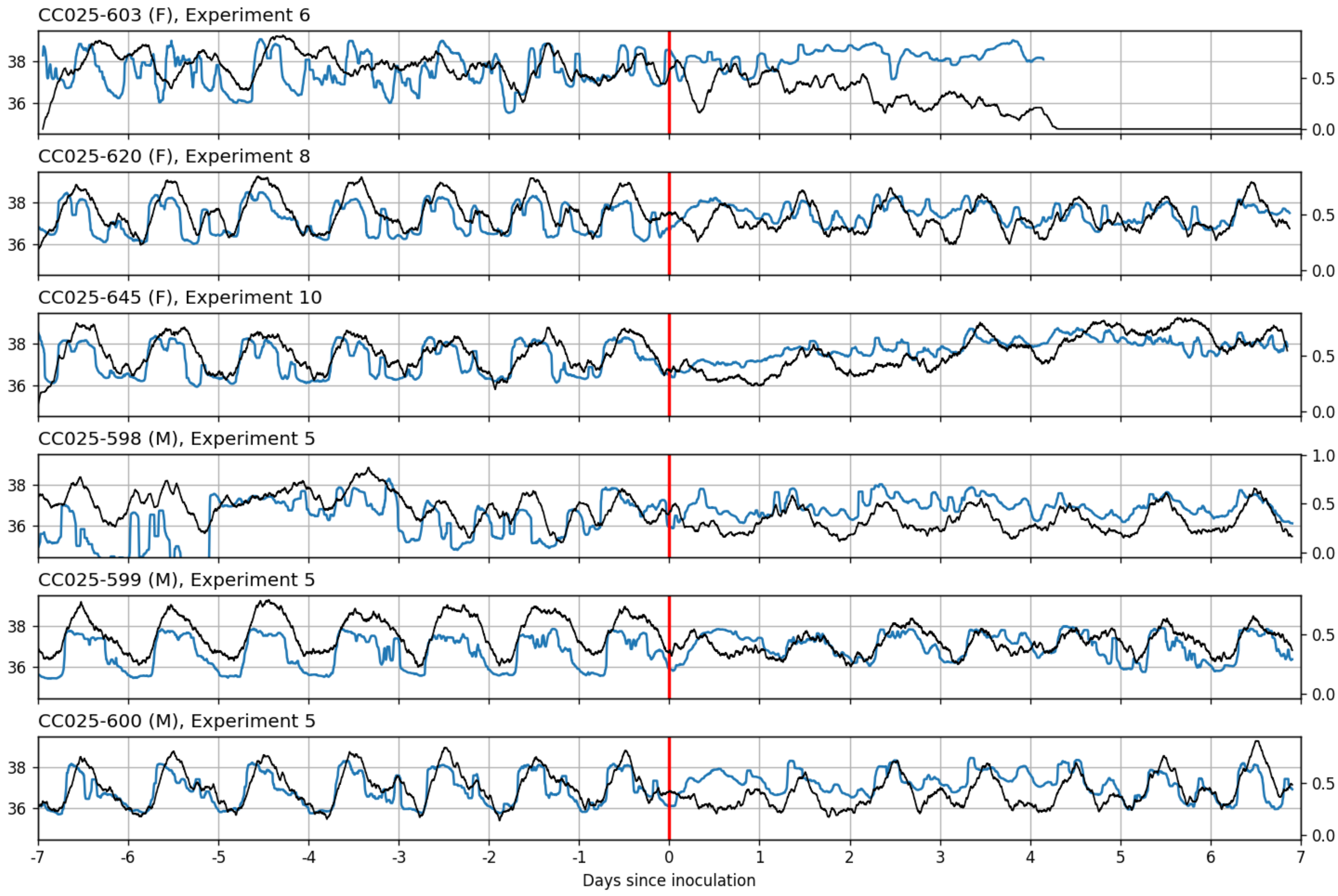

CC025-603 (F), Experiment 6
CC025-620 (F), Experiment 8
CC025-645 (F), Experiment 10
CC025-598 (M), Experiment 5
CC025-599 (M), Experiment 5
CC025-600 (M), Experiment 5
36
38
0.0
0.5
1.0
-7
-6
-5
-4
-3
-2
-1
0
1
2
3
4
5
6
7
Days since inoculation

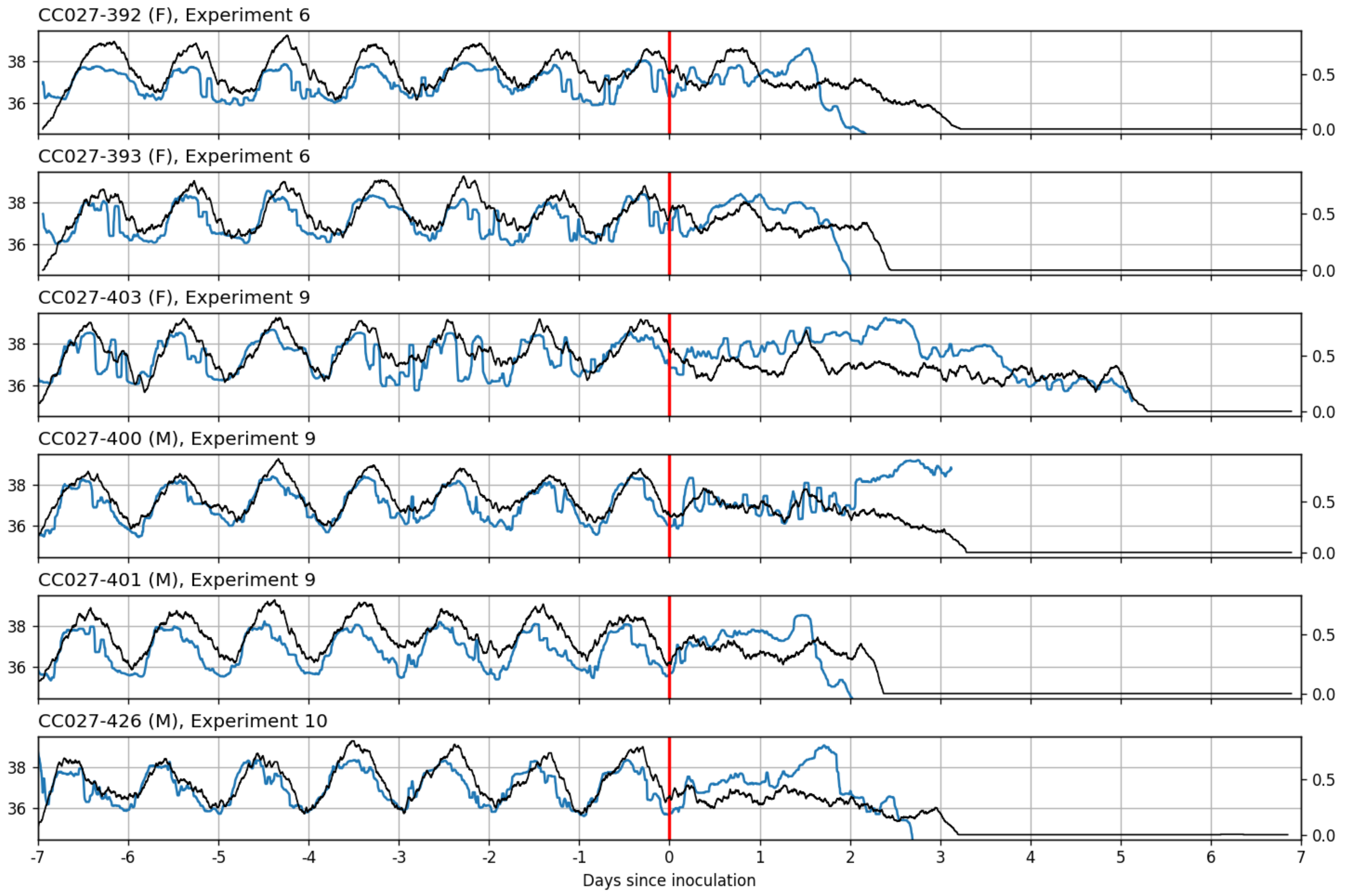

CC027-392 (F), Experiment 6
CC027-393 (F), Experiment 6
CC027-403 (F), Experiment 9
CC027-400 (M), Experiment 9
CC027-401 (M), Experiment 9
CC027-426 (M), Experiment 10
36
38
0.0
0.5
-7
-6
-5
-4
-3
-2
-1
0
1
2
3
4
5
6
7
Days since inoculation

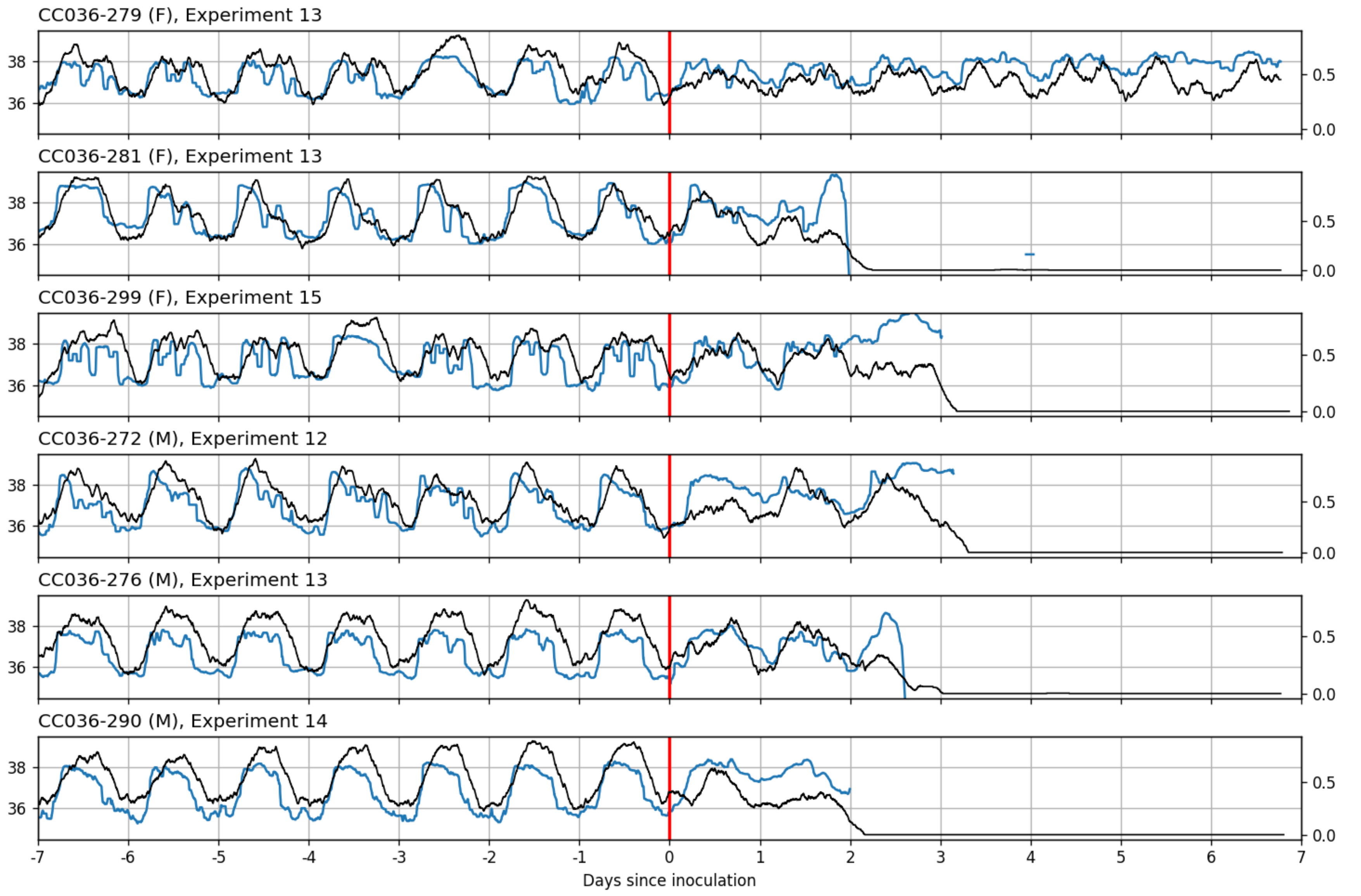

CC036-279 (F), Experiment 13
CC036-281 (F), Experiment 13
CC036-299 (F), Experiment 15
CC036-272 (M), Experiment 12
CC036-276 (M), Experiment 13
CC036-290 (M), Experiment 14
38
36
0.5
0.0
-7
-6
-5
-4
-3
-2
-1
0
1
2
3
4
5
6
7
Days since inoculation

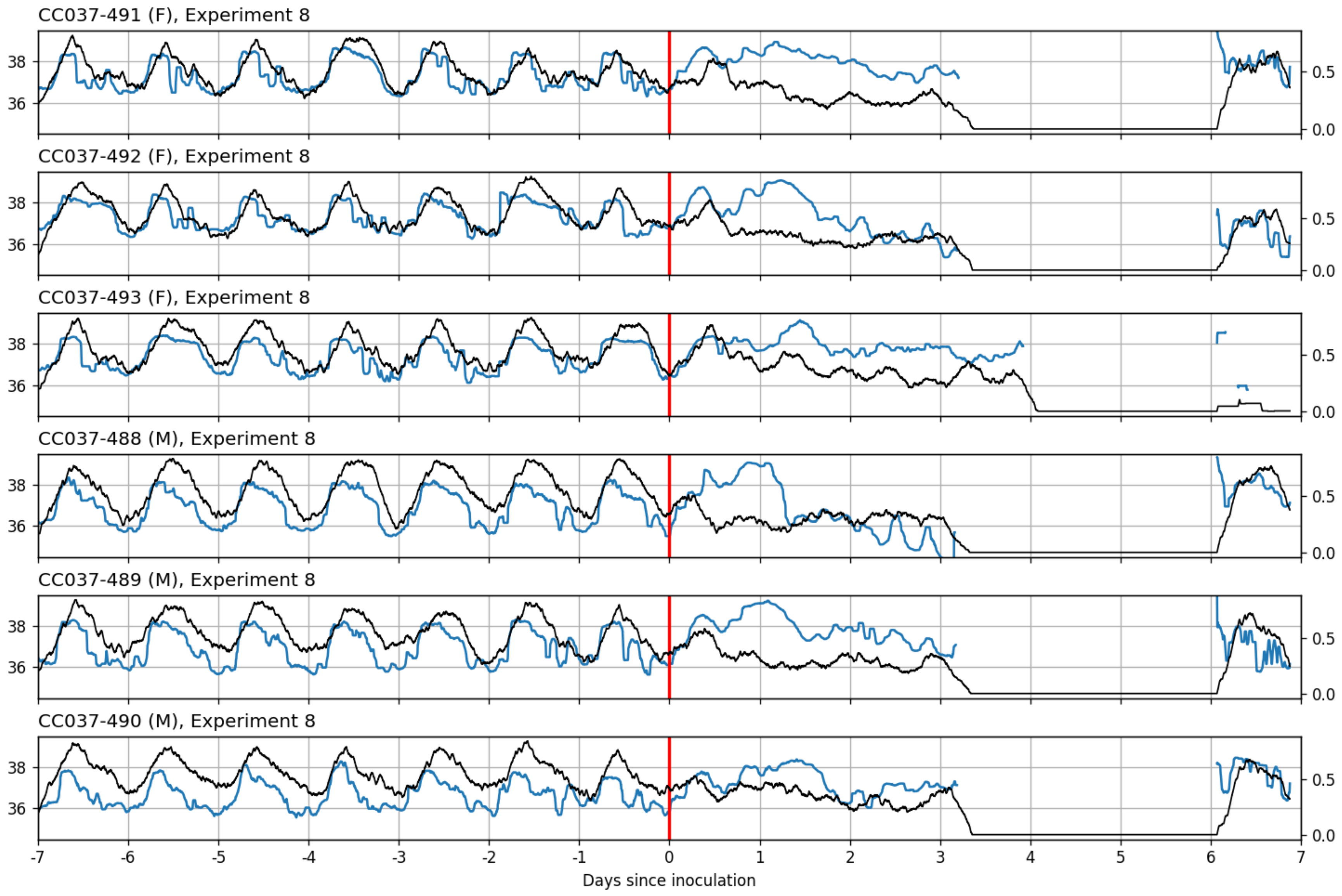

CC037-491 (F), Experiment 8
CC037-492 (F), Experiment 8
CC037-493 (F), Experiment 8
CC037-488 (M), Experiment 8
CC037-489 (M), Experiment 8
CC037-490 (M), Experiment 8
36
38
0.0
0.5
-7
-6
-5
-4
-3
-2
-1
0
1
2
3
4
5
6
7
Days since inoculation

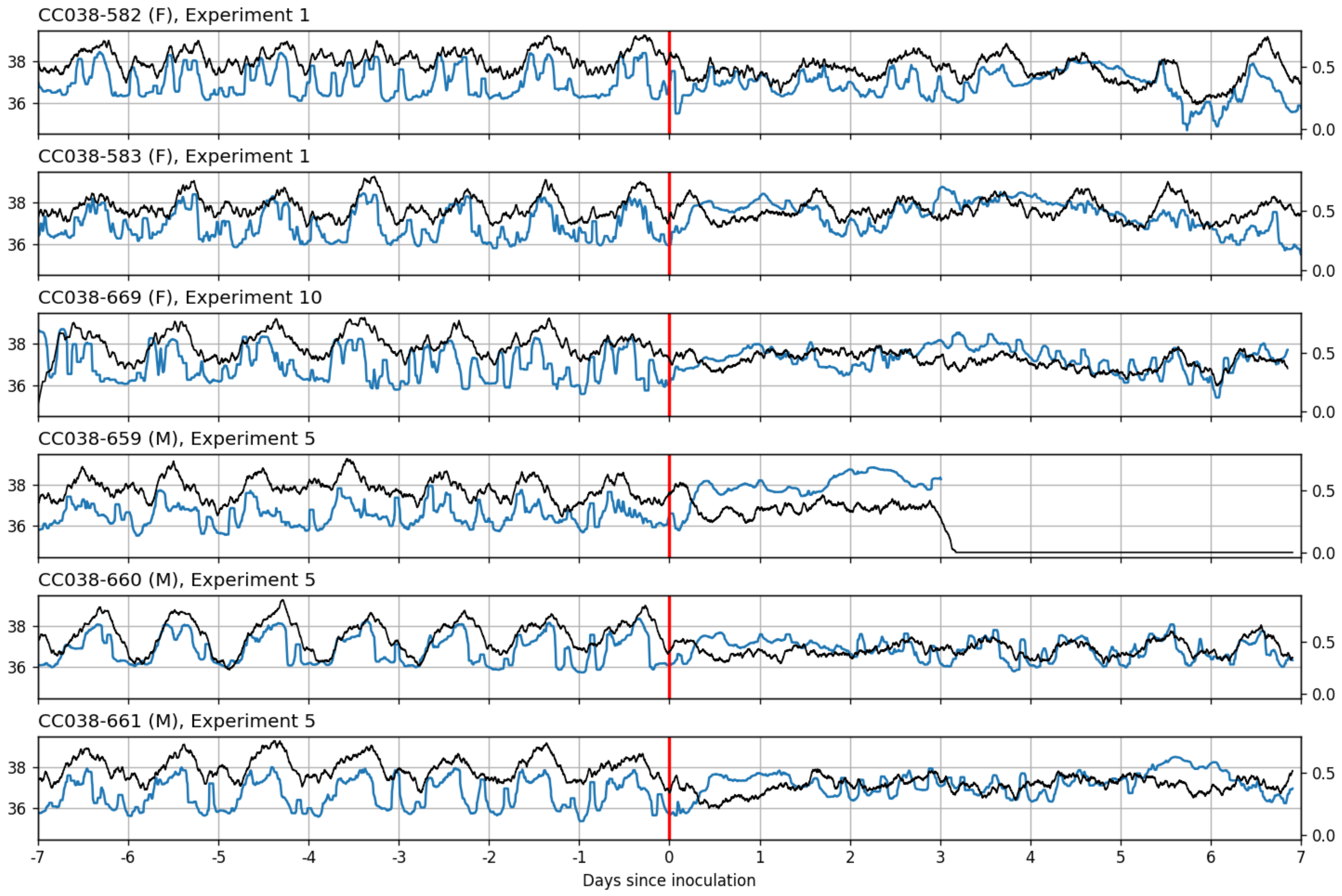
CC038-582 (F), Experiment 1
CC038-583 (F), Experiment 1
CC038-669 (F), Experiment 10
CC038-659 (M), Experiment 5
CC038-660 (M), Experiment 5
CC038-661 (M), Experiment 5
38
36
0.5
0.0
-7
-6
-5
-4
-3
-2
-1
0
1
2
3
4
5
6
7
Days since inoculation

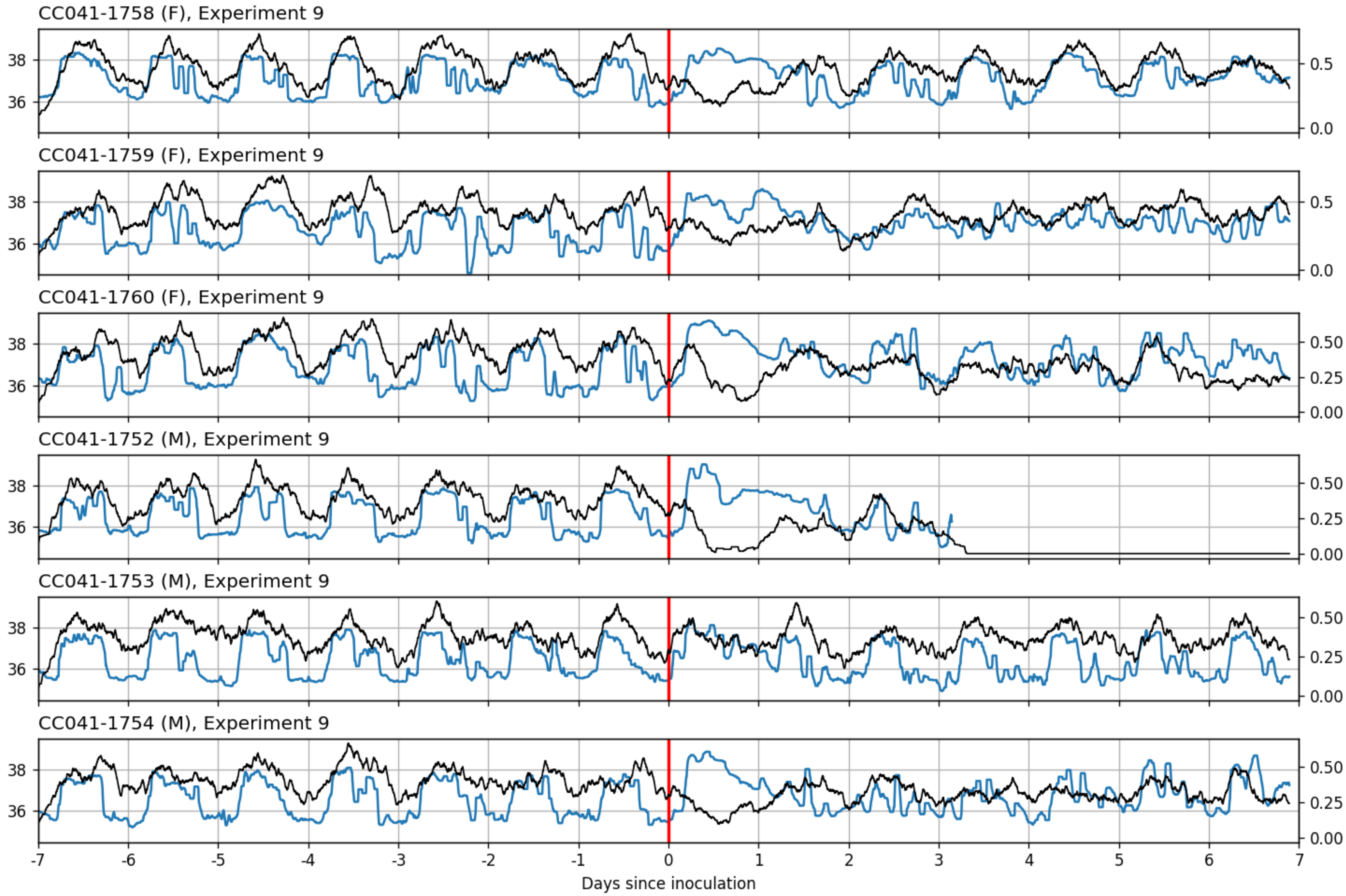

CC041-1758 (F), Experiment 9
CC041-1759 (F), Experiment 9
CC041-1760 (F), Experiment 9
CC041-1752 (M), Experiment 9
CC041-1753 (M), Experiment 9
CC041-1754 (M), Experiment 9
36
38
0.0
0.5
0.00
0.25
0.50
-7
-6
-5
-4
-3
-2
-1
0
1
2
3
4
5
6
7
Days since inoculation

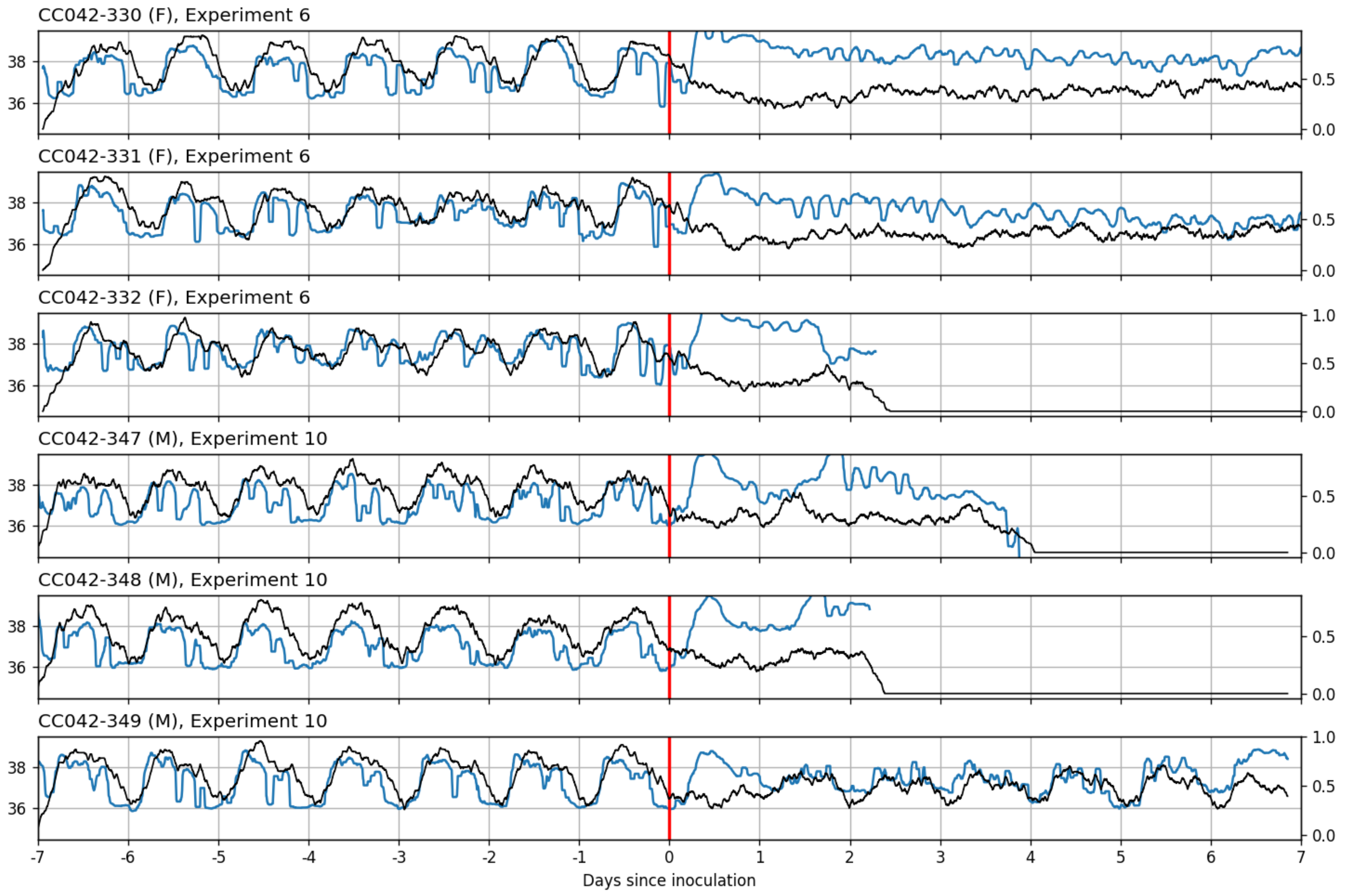

CC042-330 (F), Experiment 6
CC042-331 (F), Experiment 6
CC042-332 (F), Experiment 6
CC042-347 (M), Experiment 10
CC042-348 (M), Experiment 10
CC042-349 (M), Experiment 10
36
38
0.0
0.5
1.0
-7
-6
-5
-4
-3
-2
-1
0
1
2
3
4
5
6
7
Days since inoculation

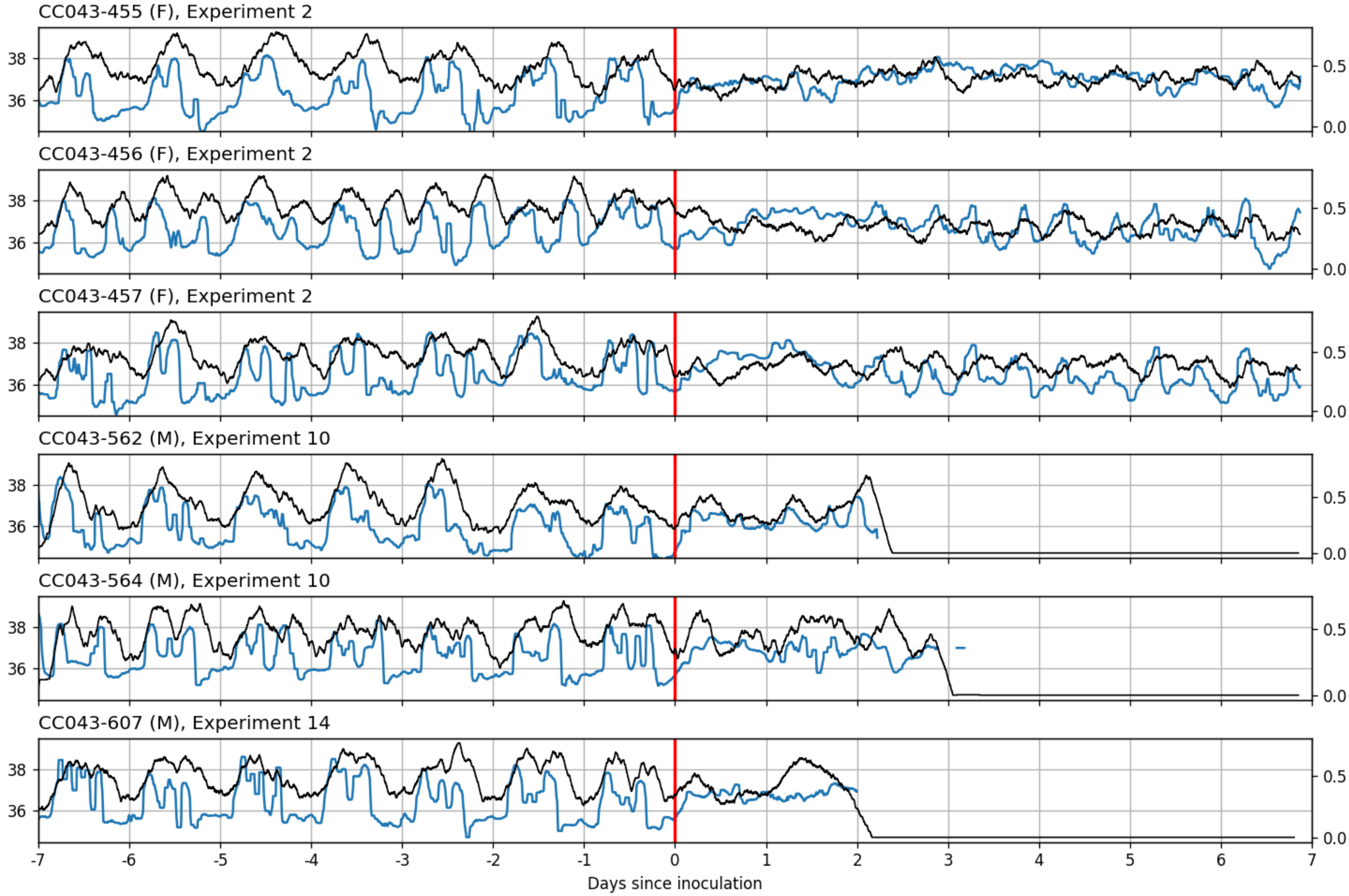

CC043-455 (F), Experiment 2
CC043-456 (F), Experiment 2
CC043-457 (F), Experiment 2
CC043-562 (M), Experiment 10
CC043-564 (M), Experiment 10
CC043-607 (M), Experiment 14
36
38
0.0
0.5
-7
-6
-5
-4
-3
-2
-1
0
1
2
3
4
5
6
7
Days since inoculation

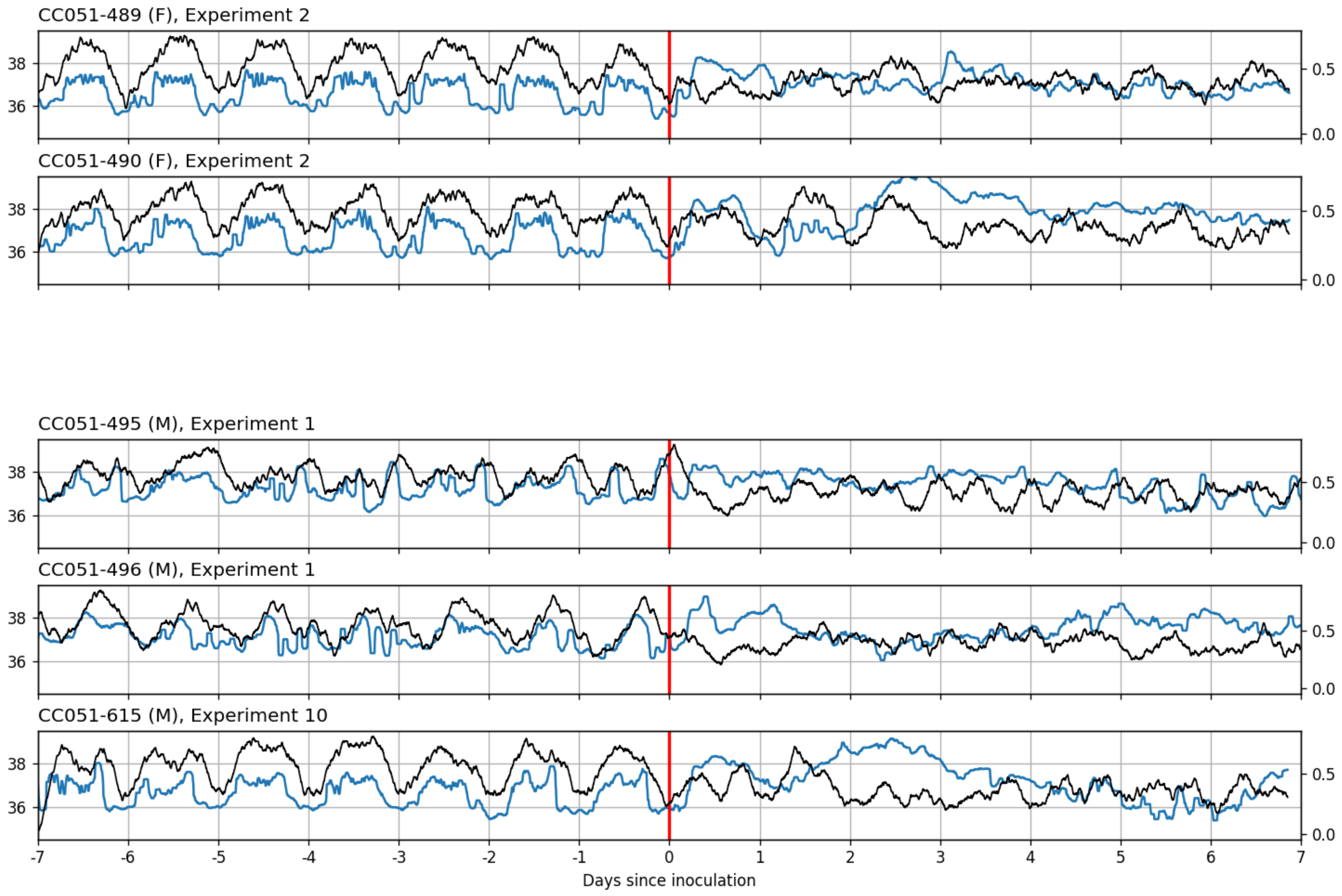
CC051-489 (F), Experiment 2
CC051-490 (F), Experiment 2
CC051-495 (M), Experiment 1
CC051-496 (M), Experiment 1
CC051-615 (M), Experiment 10
Days since inoculation

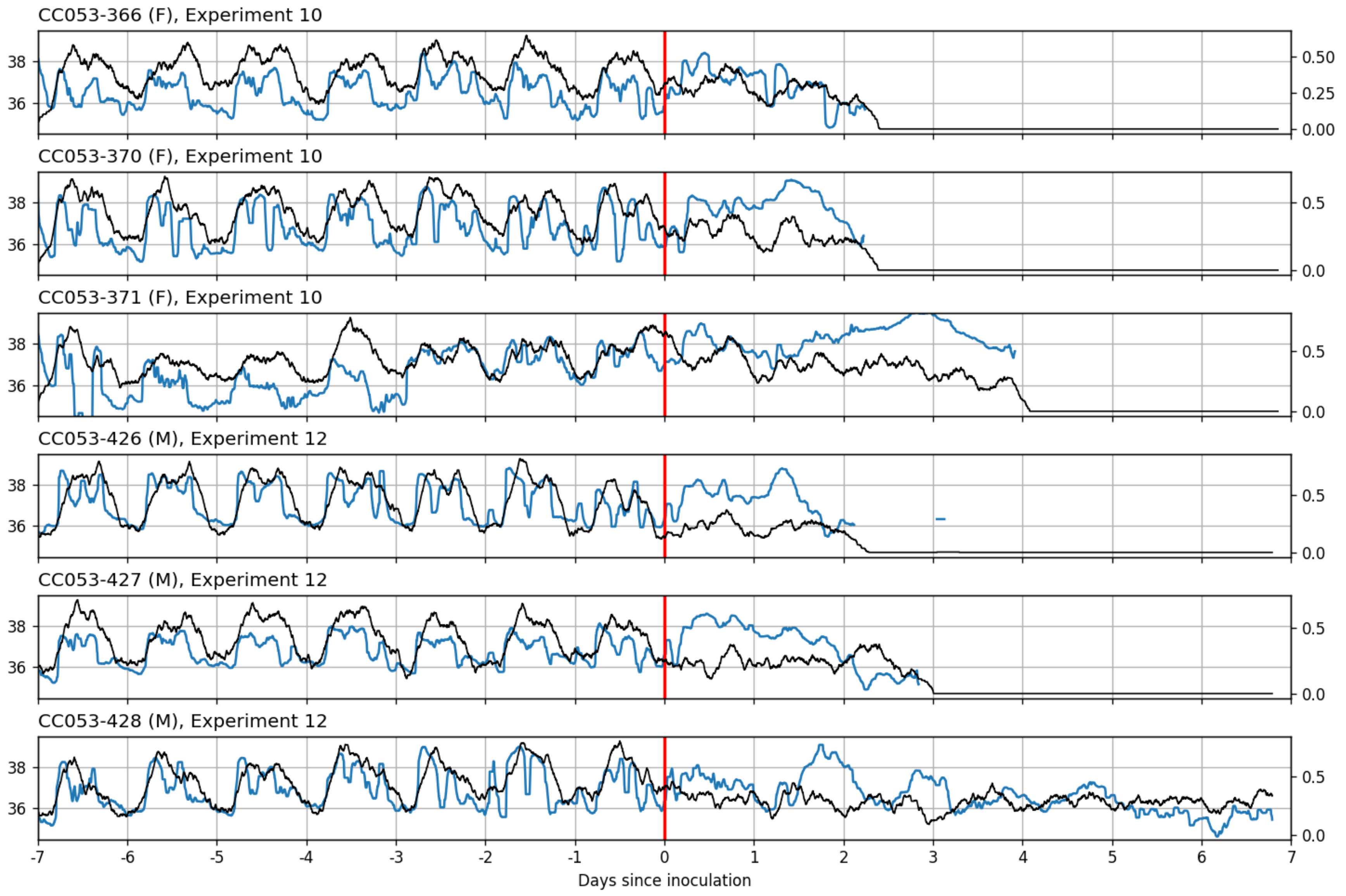

CC053-366 (F), Experiment 10
CC053-370 (F), Experiment 10
CC053-371 (F), Experiment 10
CC053-426 (M), Experiment 12
CC053-427 (M), Experiment 12
CC053-428 (M), Experiment 12
Days since inoculation

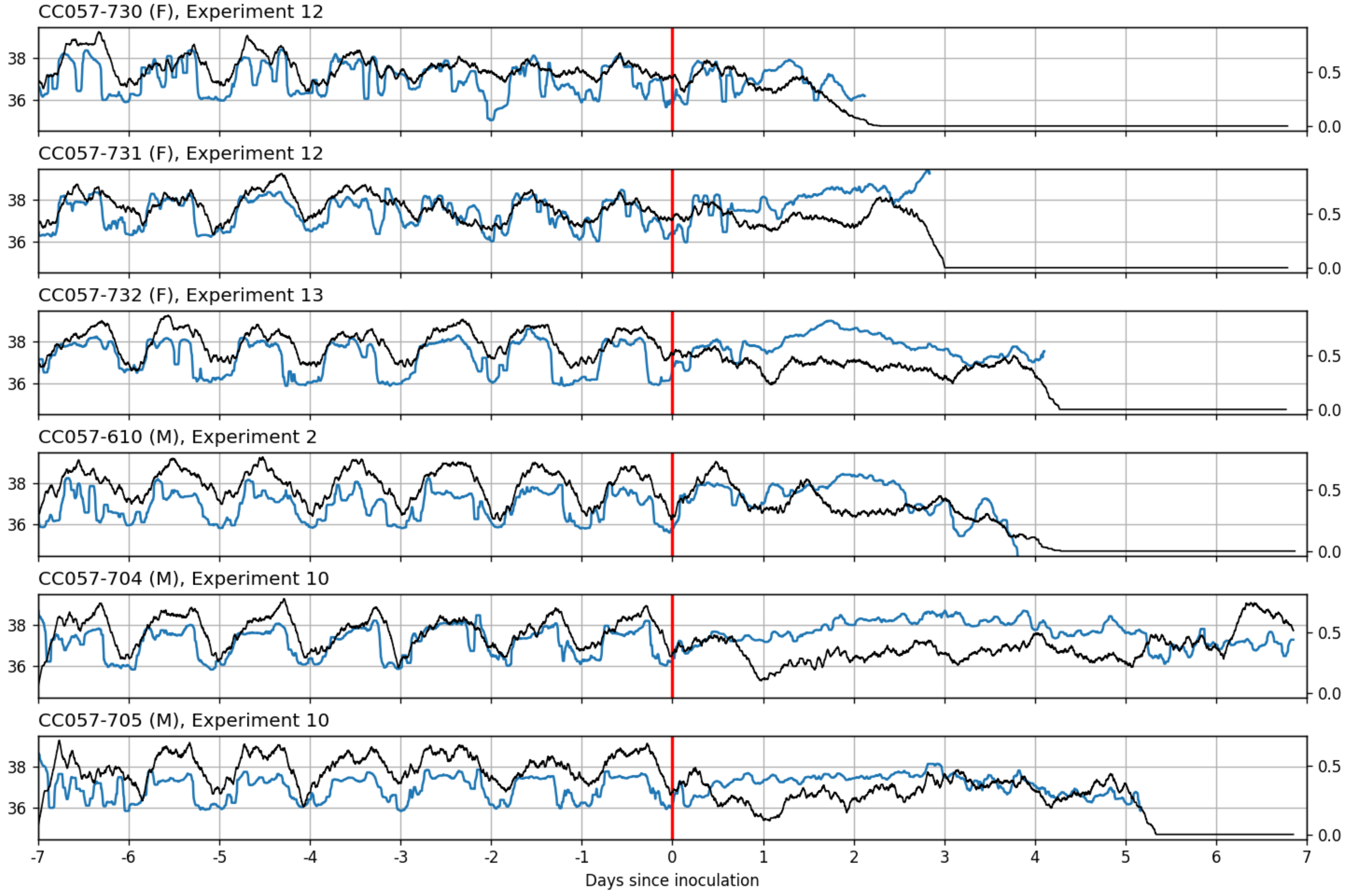

CC057-730 (F), Experiment 12
CC057-731 (F), Experiment 12
CC057-732 (F), Experiment 13
CC057-610 (M), Experiment 2
CC057-704 (M), Experiment 10
CC057-705 (M), Experiment 10
36
38
0.0
0.5
-7
-6
-5
-4
-3
-2
-1
0
1
2
3
4
5
6
7
Days since inoculation

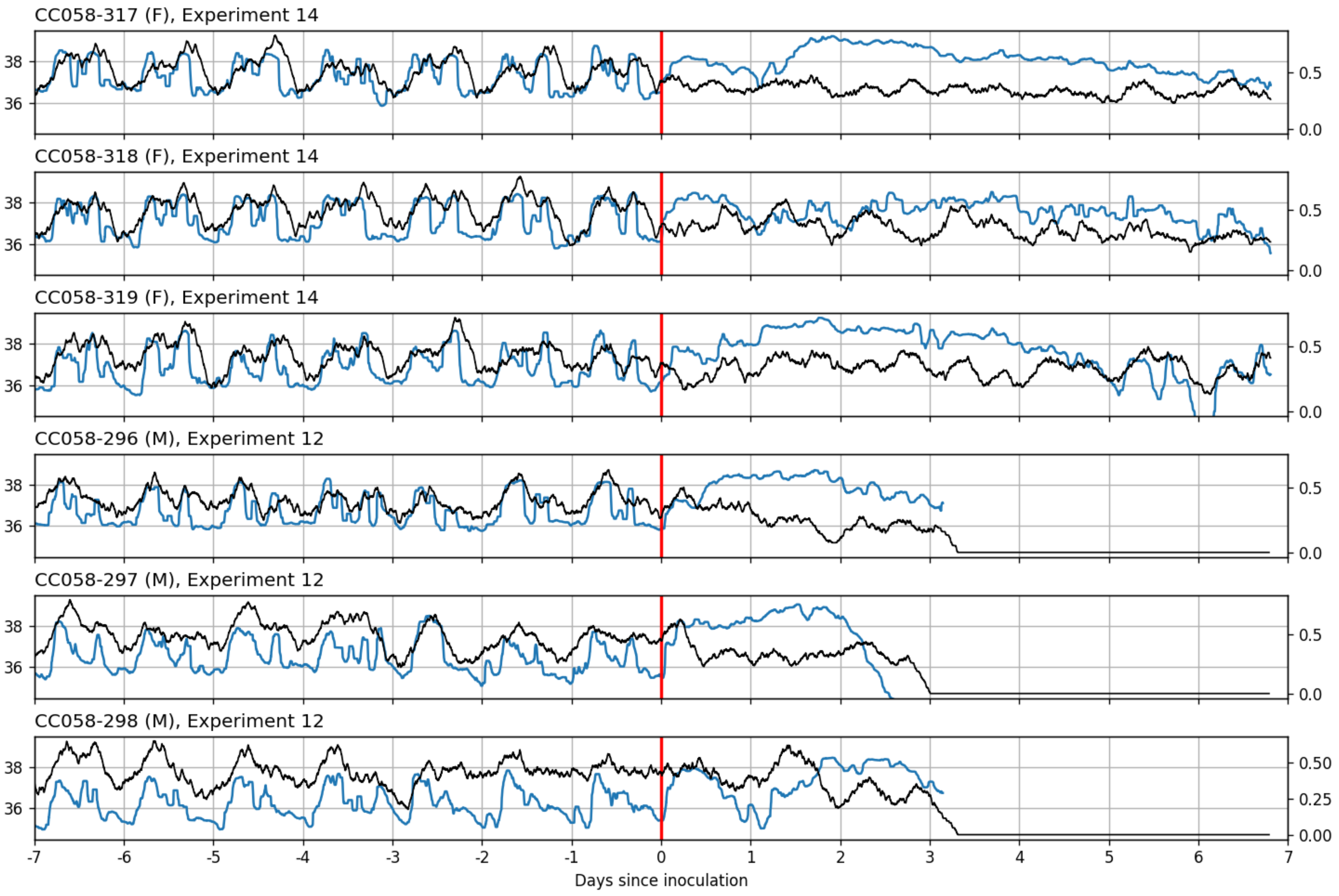

CC058-317 (F), Experiment 14
CC058-318 (F), Experiment 14
CC058-319 (F), Experiment 14
CC058-296 (M), Experiment 12
CC058-297 (M), Experiment 12
CC058-298 (M), Experiment 12
38
36
0.5
0.0
0.50
0.25
0.00
-7
-6
-5
-4
-3
-2
-1
0
1
2
3
4
5
6
7
Days since inoculation

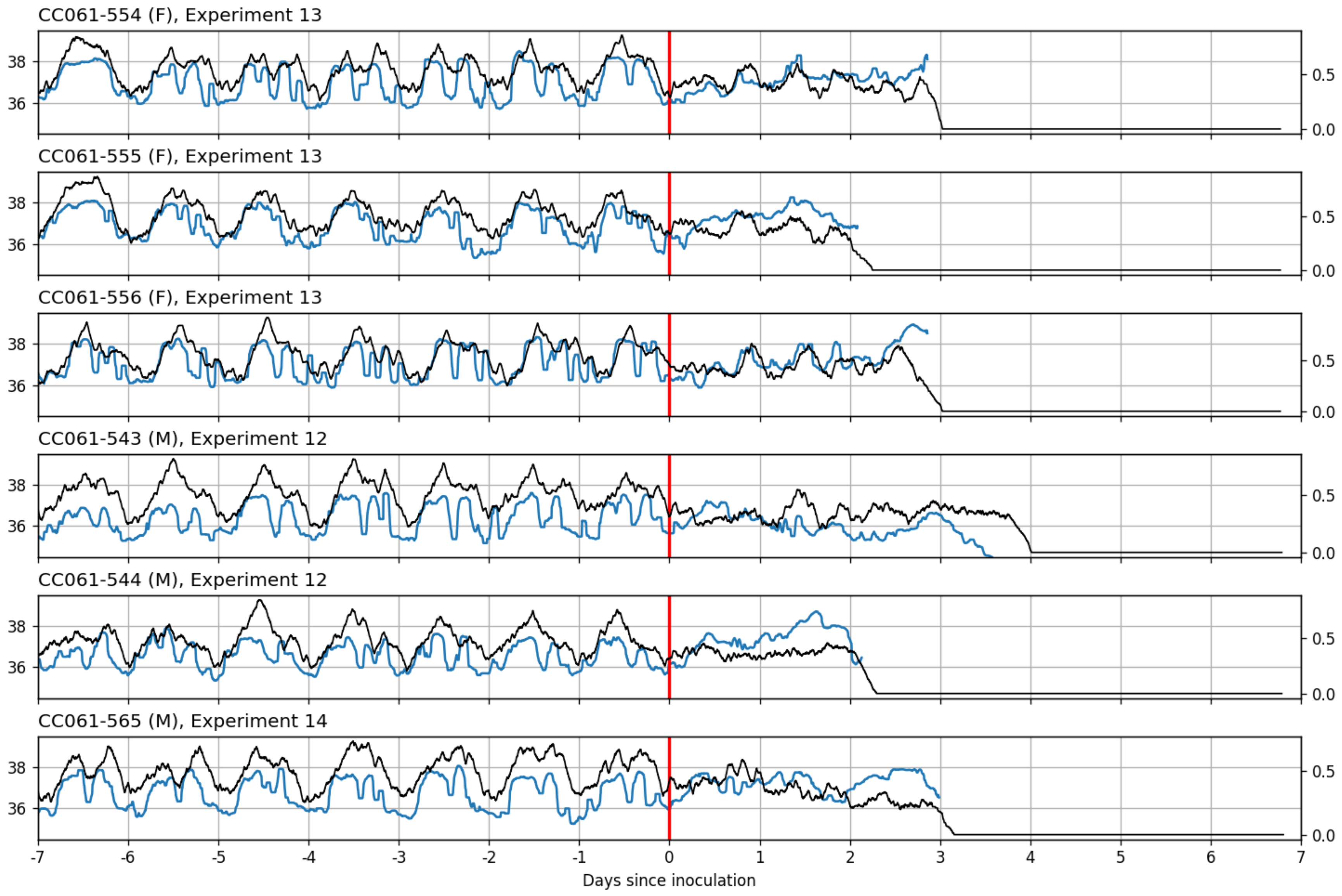
CC061-554 (F), Experiment 13
CC061-555 (F), Experiment 13
CC061-556 (F), Experiment 13
CC061-543 (M), Experiment 12
CC061-544 (M), Experiment 12
CC061-565 (M), Experiment 14
38
36
0.5
0.0
-7
-6
-5
-4
-3
-2
-1
0
1
2
3
4
5
6
7
Days since inoculation
