## Supplementary figures and images for "Collaborative Cross mice have diverse phenotypic responses to infection with Methicillin-resistant *Staphylococcus aureus* USA300"

### Fig S6

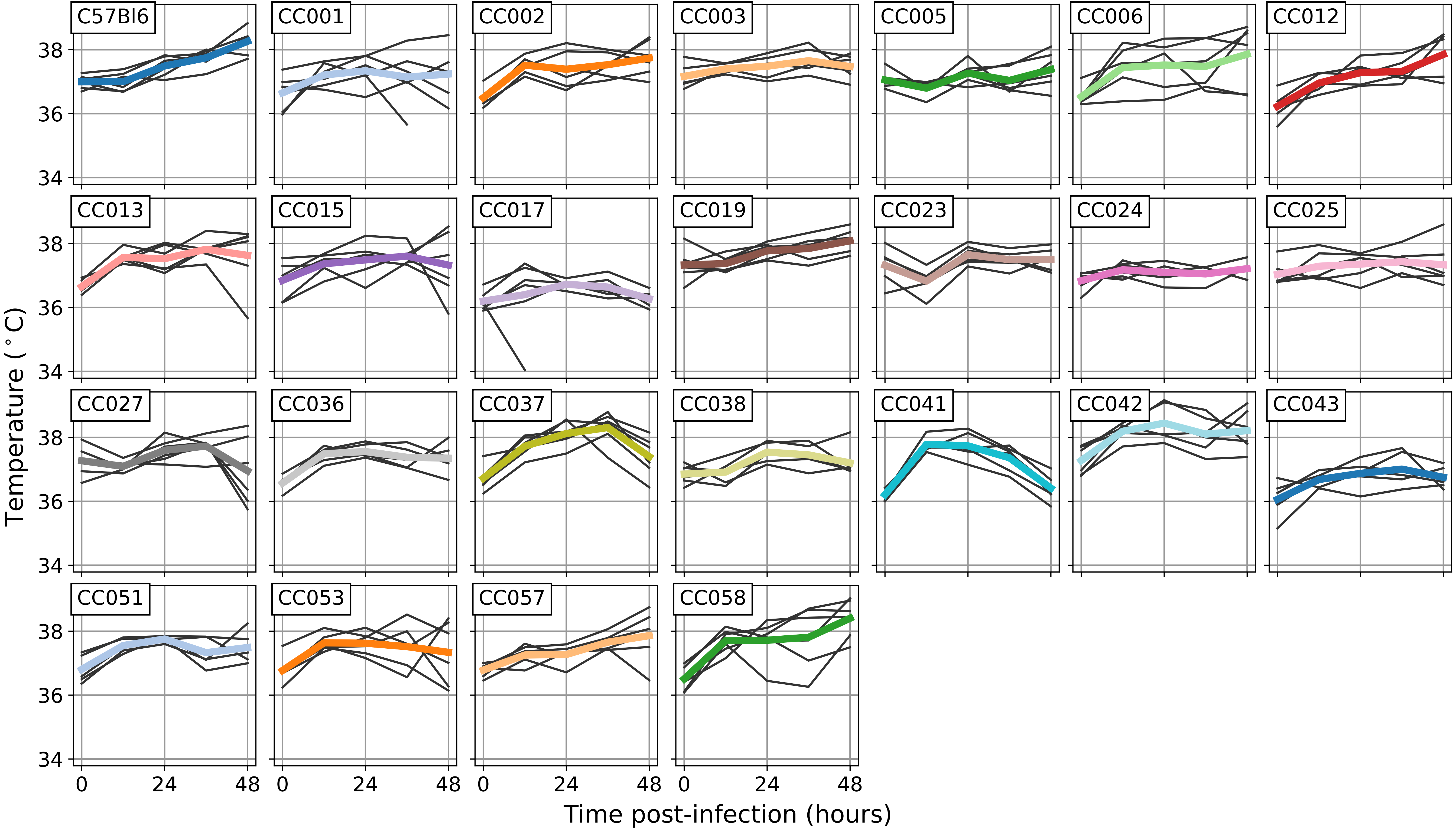
